# Validation of a molecular workflow for *Cochliomyia hominivorax* (New World screwworm) identification in field samples

**DOI:** 10.64898/2026.03.13.711645

**Authors:** Rachel Palinski, Jessica Hicks, Jeffery T. Alfred, Alec Thompson, Patrick Camp, Julia Thomas, Gleeson Murphy, Suelee Robbe-Austerman

## Abstract

*Cochliomyia hominivorax*, New world Screwworm (NWS), has become a reemerging veterinary concern in the United States due to the recent northward expansion of fly detections as far as northern Mexico. Rapid, accurate and validated detection pipelines need to be developed in the case of an incursion into the United States. Confirmatory cases are evaluated by morphological identification with no paired test to verify identifications. With the frequency of submissions of non-ideal samples, particularly from fly traps, a molecular tool would be necessary for species identification. In this manuscript, we develop and assess a pipeline including three real-time PCR assays targeting the ribosomal RNA and five sets of Sanger primers targeting the mitochondrial genome that would be used as a paired tool with morphological identification. Two of the assessed real-time PCR assays are highly specific, sensitive and repeatable requiring <1 copy per reaction for detection. Four of the five Sanger primer sets were assessed, optimized and results evaluated for potential use in preliminary geographic analysis of specimens. This workflow will expedite screening of samples, provide a method to verify results using different tools and help understand genetic variations within the mitochondria for NWS outbreaks.

## Introduction

*Cochliomyia hominivorax*, the New World screwworm (NWS), is a significant veterinary and public health concern^1,2^. With the recent rapid expansion of NWS across Central America and southern Mexico, areas from which it had been eradicated from for decades, fresh interest has blossomed to characterize the pest and reassert control over its range. Infestations of this parasite may begin with egg deposition in small wounds or body orifices. After hatching, larval stages (maggots) feed on live tissue, expanding wounds and attracting additional flies. If untreated, extensive tissue damage or even death can result. Morphological identification is the primary method for diagnosing myiasis, including for NWS; and visual methods are used to screen fly traps in high-risk areas ^3^. However, the rapid detection of the organism in low-risk areas and the identification of severely damaged specimens require additional tools, such as molecular methods, for efficient integrated surveillance. Here we describe the development of a diagnostic workflow including two species-specific and sensitive real-time PCR assays performed under universal cycling conditions. Additionally, we describe mitochondrial genome targets for Sanger sequencing conducted separately to provide confirmatory support for positive real-time PCR detections, additional species identification, and provide limited geographic information for NWS specimens. Together these assays expand available options for the rapid identification of NWS, particularly when morphological identification is challenging or not possible. The ability to link genetic variants to geographic regions may allow strengthen surveillance efforts associated with suspected incursions This manuscript provides the first step to understand genetic variants of *Coc. hominivorax*.

## Methods

### Primer and Probe Design

For real-time PCR development, both mitochondrial and ribosomal DNA (mDNA and rDNA, respectively) targets were evaluated. To identify potential mitochondrial pseudogenes on the autosomes, the mitochondria (CM118047.1) was aligned to each chromosome of GCA051144965.1 using LastZ ^4,5^. Alignment of CM118047.1 was also performed to each chromosomal assembly of *Cochliomyia macellaria* (GCA_052576185.1) with LastZ to assess pseudogene conservation between species. For rDNA analysis, the complete rDNA segment of *Calliphora vicina* (NC_088785.1:441,475-448,533) was used to identify corresponding rDNA segments in the *Coc. hominivorax* genome (GCA051144965.1) via LastZ alignment. No complete or annotated rDNA sequence for *Coc. hominivorax* was available in NCBI. Primer sets were created using Geneious Prime v 2025.2.2 (https://www.geneious.com) and the IDT PrimerQuestTool. Candidate sets were evaluated for dimerization and hairpin formation with IDT OligoAnalyzer Tool. BLAST was used to assess potential cross reaction, as were searches of raw FASTQ files with command line tools. Primer and probe sets with the most potential, *i*.*e*., not having high identity to sequences in species other than *Coc. hominivorax*, were then searched using agrep with up to three mismatches against the short-read sequence set containing *Coc. hominivorax* as well as *Coc. macellaria, Chrysomya megacephala*, and *Phormia regina*, to ensure all primers and probes were present in the *Coc. hominivorax* genomes and were not present in other species^6^. To select Sanger sequencing primers, a list of 480 published primers (Supplementary Table 1) spanning regions of the *Coc. hominivorax* genome was compiled and assessed based on the following parameters: GC content >30%, T_m_ >53°C, hairpin T_m_ <40°C, self-dimers of <5bp, homodimer ΔG >-5 kcal/mol, and heterodimer ΔG <-5 kcal/mol. Of the published primers, none either met these parameters or were bound to the preferred length limits for Sanger sequencing (1000-1400bp maximum amplicon size); therefore, we proceeded to develop new primer sets for the mitochondrial genome of *Coc. hominivorax*. An alignment of *Coc. hominivorax* sequences and one *Coc. macellaria* sequence was used to determine relevant sequencing sites. Once potential target sequences were identified, primer sets were created using Geneious Prime v 2025.2.2 and the IDT Oligoanalyzer Tool. Primer sets and probes selected appear in Table 1(Supplementary Figure 1).

**Table 1.**
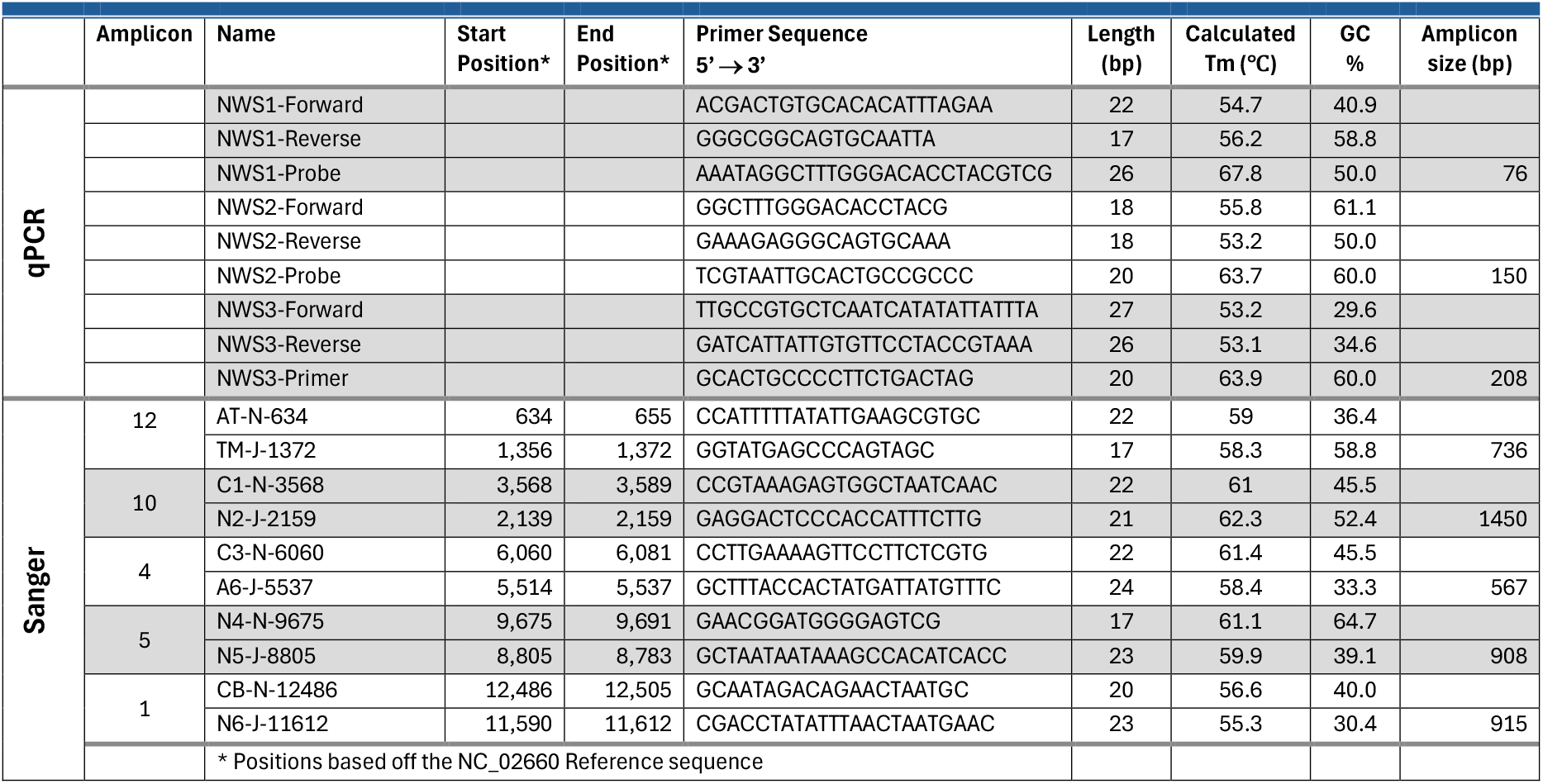
Primer and probe sequences and characteristics for the *Coc. hominivorax* real-time PCR and sequencing.

**Figure 1.**
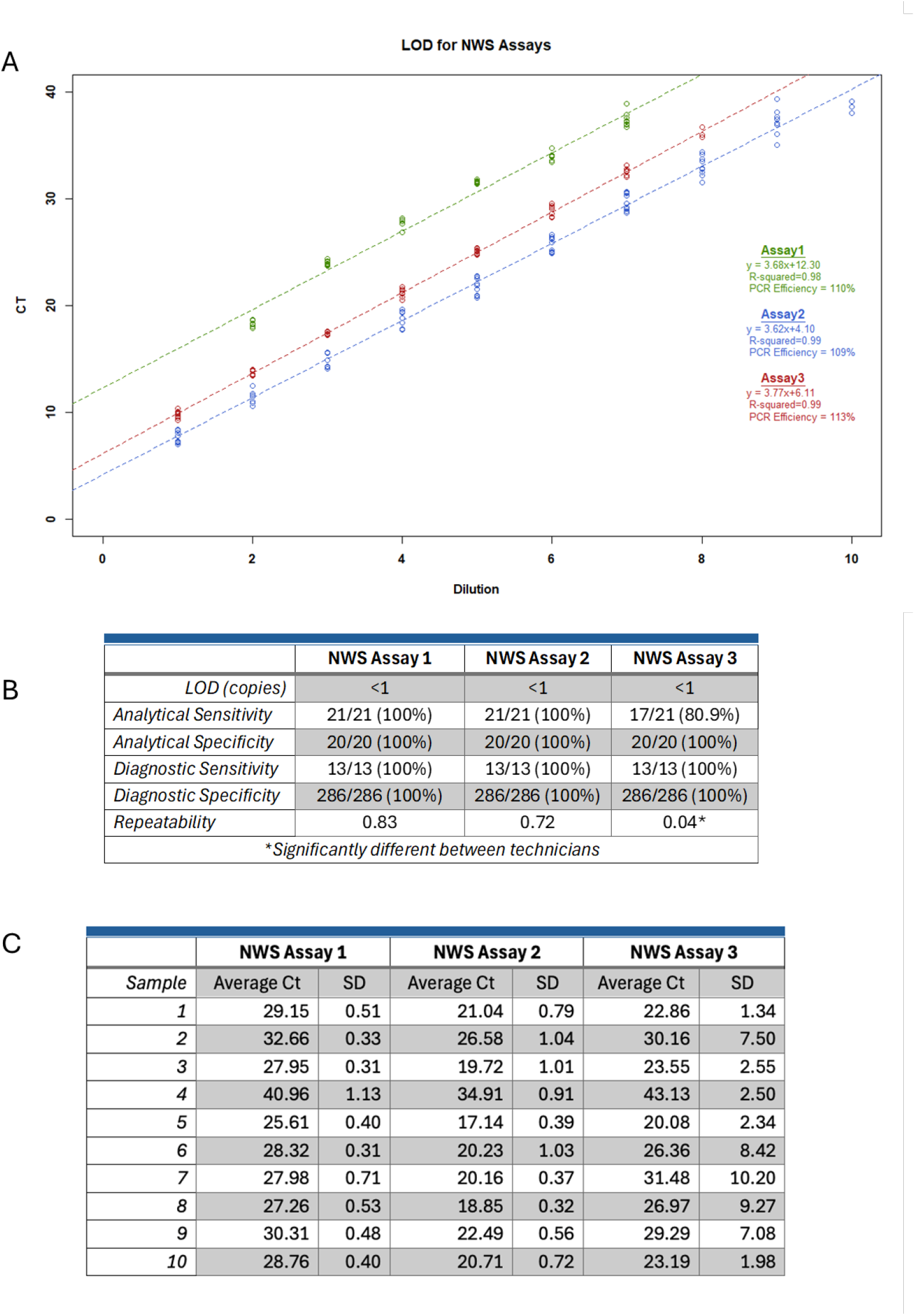
Real-time PCR results for three NWS Assays. A) LOD for NWS Assays, B) Summary of the real-time PCR results for sensitivity and specificity, and C) Summary of repeatability panel samples across three technicians. Dilutions in A are defined as 10^^-X^ where X is the dilution number displayed on the X-axis of the figure. Variance and standard deviation values above 1 are highlighted in bold in panel C. Repeatability was calculated using a one-way ANOVA in R.

### Sample source and metadata

Samples were collected as larvae from clinical myasis cases or as adult flies from environmental flytrap collections (Supplementary Table 2) as part of routine surveillance activities in each country. Larvae and a subset of fly samples were immersed in 70% ethanol and processed at the National Veterinary Services Laboratories (NVSL); the remaining fly samples were part of an ongoing flytrap surveillance effort within the United States and arrived at the NVSL with no preservative. A majority of the samples were morphologically identified before being shipped to the NVSL and all others were identified following receipt. All seven Central American countries and Mexico were represented in the sample set. The routine collection protocol of larvae for some countries includes a boiling step prior to immersing the samples in ethanol.

### Extraction

Total nucleic acid was extracted using one of three protocols: Gentra Puregene tissue kit (Qiagen), DNeasy Blood and Tissue (Qiagen) kit, or the MagMAX Core Total nucleic acid extraction kit (Applied Biosystems). In all protocols, we performed the optional digestion steps at 55°C for at least 2 hours or overnight (Supplementary Table 2). All protocols were performed based on the manufacturer’s specifications. Xeno DNA was added prior to bead beating and extraction as specified by the manufacturer.

### Real-time PCR

Real-time PCR was performed using the TaqMan™ Fast Virus 1-Step Master Mix for real-time PCR (Thermo Fisher) based on the manufacturer’s protocol with the following modifications: 1 µl of water was replaced with 1 µl of 25 mM MgCl_2_ (Millipore Sigma). The VetMAX™ Xeno™ Internal Positive Control - LIZ™ Assay was used with extracted samples and added to the PCR mastermix as specified by the manufacturer. The final concentrations of reaction components appear in Supplementary Table 3. The reactions were processed on a QuantStudio5 (Thermo Fisher) with 1 cycle of 95°C for 20 seconds, 40 cycles of 95°C for 15 seconds, and 50°C for 30 seconds with data collection (Supplementary Table 4). Annealing temperatures were tested starting at 5°C below the IDT T_m_ and further optimized as required.

To evaluate the limit of detection, a synthetic control was generated for each assay in a p-IDTSmart-AMP plasmid to be used for the dilution series. The lyophilized plasmid was resuspended in 100 µl of nuclease-free water and a dilution series created using ten 1:10 dilutions starting with the undiluted control. The dilution series was run in triplicate for three runs on each candidate real-time PCR. Copy number was calculated for the dilution series based on the following equation using concentrations of the undiluted plasmids obtained from the Qubit 4.0 High sensitivity kit according to the manufacturer’s instructions:

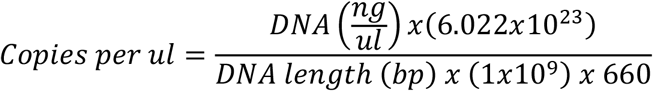

### Sanger Sequencing

The PCR amplicons were produced for Sanger sequencing using the optimized reaction and cycling conditions outlined in Supplementary Table 5. Reactions were optimized for template quantity, MgCl_2_ concentration, BSA concentration, extension time and temperature, and melting temperature. Briefly, the FastStart™ Taq DNA Polymerase, 5 U/μl kit (Roche) was used with 1.5 mM MgCl_2_, 1.25 mM DMSO, and 2 µl of template diluted 1:4 (if applicable) in nuclease-free water. DNA was only diluted 1:4 if the concentration obtained on the Qubit was above 20 ng/µl. Cycling conditions can be found in Supplementary tables 4 and 5. Amplicons were purified with ExoSAP-IT ™ Express PCR product Cleaner (Applied Biosystems) and prepped for sequencing using BigDye™ Terminator v3.1 Cycle Sequencing Kit. Unincorporated dyes were removed using BigDye Xterminator ™ Purification Kit (Thermofisher). The sequencing preps were run on an Applied Biosystems™ SeqStudio™ 24 Flex Genetic Analyzer. Reads were analyzed and aligned in Geneious 2025.2.2. SNPs were called using the ‘Find Variations/SNPs’ function and confirmed visually once called.

### Whole Genome Sequencing

DNA extracted from whole flies or maggots was sequenced on an Illumina NextSeq 2000 with P3 300-cycle flow cells. Library preparation was performed using Illumina DNA Prep. Output reads for each isolate were aligned to assembly GCA051144965.1 with vSNP Step1 where read quality criteria were gathered with SeqKit, and alignment metrics were obtained for the whole genome. Individual elements of specific interest, specifically the mitochondria and rDNA segments, were further inspected with this short-read data.

### Bioinformatics, including creating a tree of sequences

Sanger amplicons were aligned with MAFFT and then merged in Geneious Prime v 2025.2.2. Model selection was performed with Modeltest-NG, and a phylogenetic tree was created in RAxML with 1000 bootstraps based on the GTRCAT model.

### Statistical Analysis

Statistical calculations were determined using R version 4.3.0 and R Studio 2023.03.0^7^. For the repeatability analysis, a model was created using the base functions and a summary for one-way ANOVA for all assays.

## Results

### Real-time PCR Limit of Detection (LOD) and Analytical Sensitivity

The control plasmid for each assay was diluted from 10^-1^ to 10^-11^ and run in triplicate for three runs. The corresponding Cts were assessed for linearity resulting in an R^2^ of 0.98 for Assay 1, 0.99 for Assay 2, and 0.99 for Assay 3. Within the plates, the R^2^ values were 0.99 across all plates.

The LOD was determined at the last dilution in which all replicates were detected; therefore, the LOD for Assays 1, 2, and 3 were calculated to be 10^-7^, 10^-9^, and 10^-7^, respectively, each corresponding to <1 copy per reaction. Figure 1A displays the dilution series results for each assay across the three runs. The 10^-1^ dilution was removed from Assay 1 due to detection inconsistency.

Additionally, 21 *Coc. hominivorax* maggots were tested using all real-time PCR assays, collected from the following eight countries: Belize (2), Panama (3), Costa Rica (3), Nicaragua (3), Honduras (3), El Salvador (3), Guatemala (3), and Mexico (1). Of these, Assays 1 and 2 correctly identified all samples as *Coc. hominivorax* with Cts ranging from 18.7 to 36.8 and 14.5 to 37.3, respectively. Assay 3 correctly identified 17/21 samples with Cts ranging from 17.4 to 31.9. This resulted in an analytical sensitivity of 100% for Assays 1 and 2 and 80.9% for Assay 3 (Figure 1B). All samples were sequenced using the Sanger amplicon primers.

### Real-time PCR Analytical Specificity

A total of 20 flies from 5 closely related species [*Compsomyiops callipes* (4), *Phormia regina* (4), *Chrysomya rufifacies* (4), *Cochliomyia macellaria* (4), *Chrysomya megacephala* (4)] were tested on all assays to assess analytical specificity. None of the closely related species yielded detectable Ct values across any of the three assays resulting in analytical specificities of 100% for all assays. (Figure 1B)

### Diagnostic Sensitivity and Specificity

Diagnostic sensitivity and specificity were assessed by individually testing 299 fly legs extracted using the MagMAX Core nucleic acid kit based on the manufacturer’s specifications for tissue digestion. These samples were blinded to the technician and results were assessed following the run completion.

A total of 13/299 samples had been identified *a priori* as *Coc. hominivorax* by morphological identification. Of these, all assays correctly identified the *Coc. hominivorax* samples with Cts ranging from 30.67-32.47 (Assay 1), 20.17-24.96 (Assay 2), and 20.06-27.76 (Assay 3). Two hundred and eighty-six samples morphologically identified *a priori* as non-*Coc. hominivorax* were tested across all three assays; and all three assays correctly classified these samples as such with no Ct determined. Internal exogenous control Xeno values ranged from 33-37. The assay was repeated if Xeno values rose above a Ct of 37. This upper limit was calculated by identifying 2 standard deviations above the average Ct across all samples from a single assay. These assessments resulted in diagnostic sensitivities and specificities of 100% for all three assays. (Figure 1B).

### Repeatability

Assay repeatability was tested with a panel of 10 samples of varying Ct values (17-36). The samples were tested by three technicians blindly and in duplicates. For Assay 1, all samples except 1 had a variance and standard deviation below 1 (Figure 1C). Assay 2 had all but 3 samples with variances and standard deviations below 1. Assay 3 had highly variable Cts with variances between 0.98 and 20.8 and standard deviations between 0.99 and 4.5 across all replicates and samples. Average Cts across all technicians and replicates were between 6 and 8.5 Cts lower in Assay 2 than Assay 1 in the repeatability runs, while Assay 3 had Cts between 0.6 and 6.3 higher than Assay 2 across the runs. Between Assay 1 and 3, the Cts were largely lower in Assay 3 by between 3.2 and 6.29 Cts. Of the three assays, Assays 1 and 2 were by far the most repeatable; however, Assay 2 resulted in 3-6 Cts lower for the same samples across all runs. A one-way ANOVA test for each assay generated P-values of 0.83 for Assay 1, 0.72 for Assay 2, and 0.04 for Assay 3; so, with significance set at p<0.05, Assay 3 did not meet the repeatability criteria (Figure 1C).

### Bulk Fly leg Samples

Four samples were extracted that each included 5-10 non-*Coc. hominivorax* legs and one *Coc. hominivorax* leg and tested across all three real-time PCR assays. Each assay correctly assessed the presence of *Coc. hominivorax* with Cts ranging from 31-39, 20-29, and 24-30 for Assays 1, 2, and 3, respectively. These samples were used with the Sanger primers to amplify target regions. The resulting sequences correctly identified all specimens in the samples.

### Sanger Sequencing

Samples from each of the aforementioned eight countries were subjected to Sanger sequencing using the primer sets identified in Table 1. Additionally, samples that were positive in the diagnostic sensitivity and specificity analysis were also subjected to Sanger sequencing. All primer sets amplified the target consistently with the PCR results for Assay 2. Sanger primers were also assessed for specificity using the same sample set run on the real-time PCR identifying three of the primer sets that cross-reacted with at least one non-*Coc. hominivorax* species. Figure 2A portrays a high degree of variability between the *Coc. hominivorax* sequences from countries such as Panama and Honduras, whereas no variability was demonstrable amongst Costa Rica, Nicaragua, Guatemala, and El Salvador. An assessment of broodstock samples from the sterile fly production facility in Panama (COPEG) (Figure 2B) displays moderate variability between sequences from the same genomic regions. Additionally, the sequences of two known sterile and two known fertile flies were assessed (data not shown). Between these Sanger sequences, no differential SNPs were identified in the regions amplified, suggesting no DNA mutations in the consensus of the mitochondrial genome were caused by the radiation-related sterility, however, the sterility status was unknown for the bloodstock sequences in Figure 2B and therefore needs to be evaluated further.

**Figure 2.**
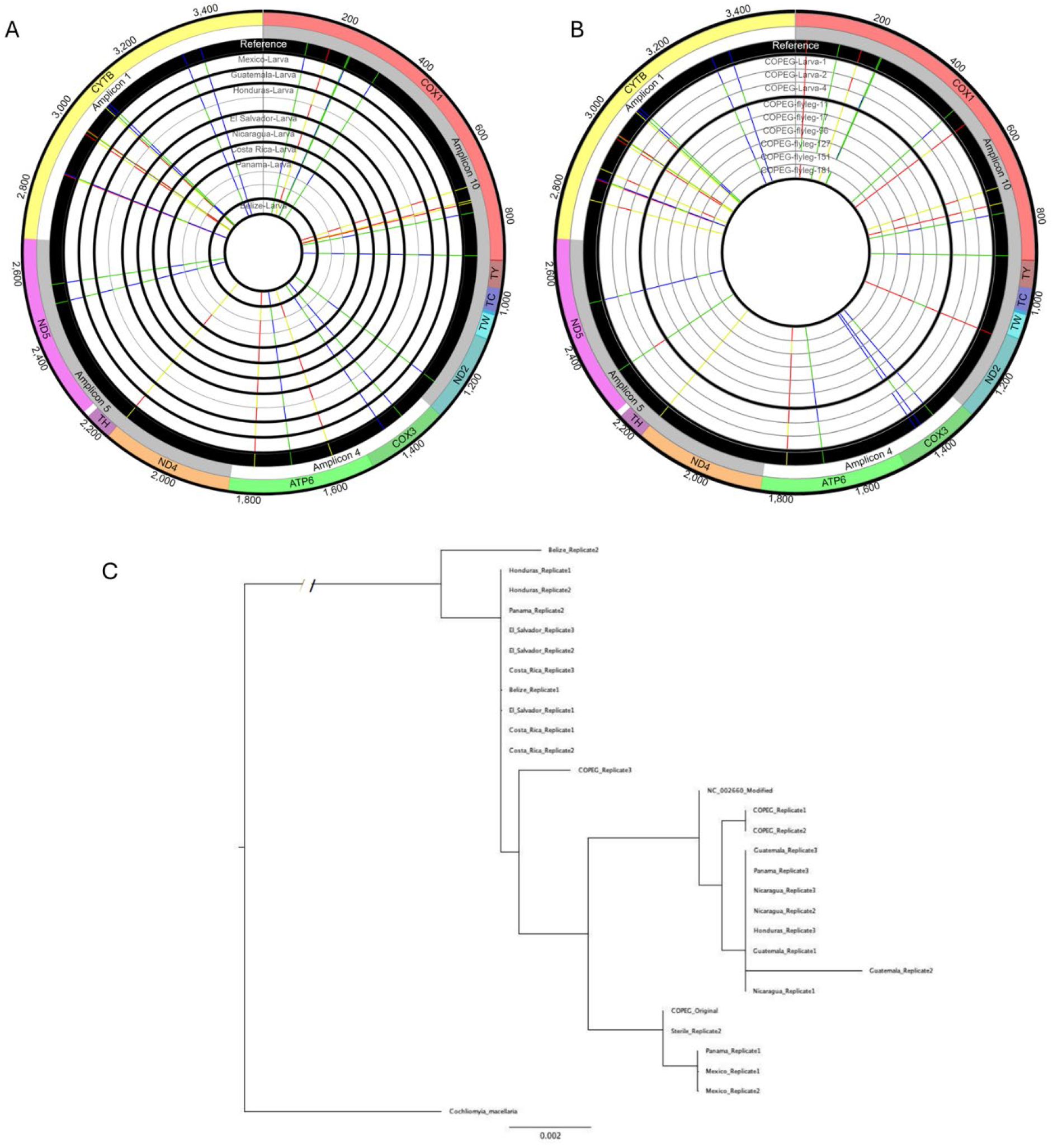
Sanger sequence results from *Coc. hominivorax* samples. A) Shown are Sanger sequences collected from samples from various countries as specified in the figure. If the sequences from these countries were identical, only one sequence is displayed in the figure. B) Displays sequences collected from flies from the sterile fly production facility in Panama. C) Tree with 1000 bootstraps of the concatenated sequences from this study. All SNP sites are defined based on the NC_002660 reference. Sequences that were identical were not included.

Concatenated sequences from the five Sanger primer sets were used to generate a phylogenetic tree. From this analysis, two supported clades appear: the first consists of Belize, Costa Rica, and El Salvador, and the second clade includes the countries Guatemala, Mexico and Nicaragua. Isolates from Honduras and Panama occur in both clades.

### Primer probe design

Mitochondrial targets were initially attempted; however, the presence of pseudogenes and lack of SNP accumulation in potential target regions resulted in primers with less discrimination between species than potential rDNA targets (data not shown).

Alignment of the rDNA segment from *Ca. vicina* only occurred on the X (CM118045.1) and Y (CM118046.1) chromosomes with 7 and 5 tandemly repeated copies, respectively. Evaluation of the rDNA in the short-read alignments revealed an approximately 600bp region from the 3’ end of the 18S gene and extending into ITS1 region that appeared unique to *Coc. hominivorax* and was not observed in other species in the data set. This region was used for real-time PCR primer and probe design. While some of the final primers and probes from this region were observed in the raw-read search, the entire set was never observed in any species except *Coc. hominivorax*.

## Discussion

Historically, restriction fragment length polymorphism (RFLP) analyses have been the standard to detect *Coc. hominivorax* in unknown samples; however, with the recent outbreak in Latin American countries, additional emphasis has been placed on developing modern molecular approaches to detect and characterize *Coc. hominivorax* in diagnostic samples ^8–10^. Random amplified polymorphic DNA (RAPD) conventional PCR techniques have also been assessed to target *Coc. hominivorax* ^11–13^. This technique uses 10-mer primer sets and assesses the amplicon sizes to differentiate species. This technique has also shown some promise to identify differences in regionality of *Coc. hominivorax* samples utilizing an amplicon size of approximately 860bp for differentiation^14,15^. Likewise, a commercial real-time PCR kit is available with a similar amplicon size and is marketed for *Coc. hominivorax* detection. To our knowledge, no data is publicly available assessing the validation of this commercial kit. Furthermore, the sample size for testing was limited for the RAPD-PCR assays. Additional work is required to compare the assays to the assays in this paper.

To date, there is one other real-time PCR assay described in the literature which differentiates *Coc. hominivorax* from closely related species using a mitochondrial target ^16^. However, that assay had some limitations including a lower specificity (95.8%) with higher Ct values in positive samples (25-35), as compared to the higher specificity and higher Ct values reported in our work. Our manuscript assessed three real-time PCR assays to differentiate *Coc. hominivorax* from other closely related fly species by targeting regions of the rDNA sequence unique to *Coc. hominivorax*. These regions are not known to have pseudogenes, and are therefore preferable for detection. In contrast, the mitochondrial targets described the previously published assay, contain pseudogenes, which can be found throughout the genome and across many species with high homology to functional genes, and can cause false positives for real-time PCR. Of the assessed assays, two of the three specifically targeted *Coc. hominivorax* in all samples tested resulting in an analytical and diagnostic sensitivity and specificity of 100% with Ct values as low as 21 for individual fly legs. Additionally, these assays were tested for repeatability and positive results confirmed by a set of Sanger primers targeting the mitochondrial genome for all positive samples. Furthermore, we validated the analytical sensitivity and specificity of our PCR assays against *Coc. hominivorax* from other countries, the breadth of which has not been performed previously.

The genome of *Coc. hominivorax* is a complex eukaryotic genome putatively composed of 5 chromosomes and 82 unplaced scaffolds encompassing 455Mb^17^. As the annotations and assembly are not complete for this organism, molecular assays are difficult to develop. Additionally, the genome is highly repetitive and includes pseudogenes that can prevent the identification of adequate molecular targets^18^. While the GC content has not been determined for the *Coc. hominivorax* complete genome, the mitochondrial GC content is about 23% making assay development a challenge and requiring modified PCR protocols to amplify^19^. In this study, we assessed hundreds of published primers and internally developed primers against stringent *in silico* standards (GC content >30%, T_m_ >53°C, hairpin T_m_ <40°C, self-dimers of <5bp, homodimer ΔG >-5 kcal/mol, and heterodimer ΔG <-5 kcal/mol) resulting in the development of three real-time PCR assays and five Sanger primer sets. Of these, only two real-time PCR assays and four Sanger primer sets met criteria required for diagnostic testing.

The samples used in this study were collected with no preservative or immersed in 70% ethanol either directly or following a boiling step^1^. Since these methods are routinely utilized in the field to collect *Coc. hominivorax* samples, we ensured that our assays were able to detect and amplify DNA sequences from such samples. The samples collected from fly traps are biased toward *Cochliomyia* spp. and carrion flies due to the use of a specialized lure^20^ and an initial morphological screening in the field for *Cochliomyia* spp. prior to submission; therefore, additional diagnostic sensitivity and specificity analyses may be required to confirm the applicability of our assays in the context of a broader range of fly species. Further validation is required for *Coc. hominivorax* detection in bulk fly samples to optimize the pool size; however, in the pools of up to 10 that were tested, the assays identified *Coc. hominivorax* with 100% sensitivity with Cts of 31-39, 20-29, and 24-30 for Assays 1, 2, and 3, respectively. Thus, this pool size may already be appropriate for fly trap use. Our results are similar to the previously published real-time PCR results for bulk fly samples with the exception that the Cts were much lower for Assay 2 than the published assay^21^.

This work also provides a workflow for testing individual fly leg, maggot, and whole fly samples for *Coc. hominivorax* detection and provides the groundwork for phylogeographic analyses (Figure 3). Further studies will assess the utility and validation in bulk fly trap samples. Furthermore, the generated workflow was tested and validated in this manuscript to be used for *Coc. hominivorax* testing. The proposed workflow is also highly modifiable to other species and could identify non-*Coc. hominivorax* species such as *Coc. macellaria, Ch. rufifacies, P. regina, Com. callipes*, and *Ch. megacephala* by modifying the primers in the target regions to detect these species.

**Figure 3.**
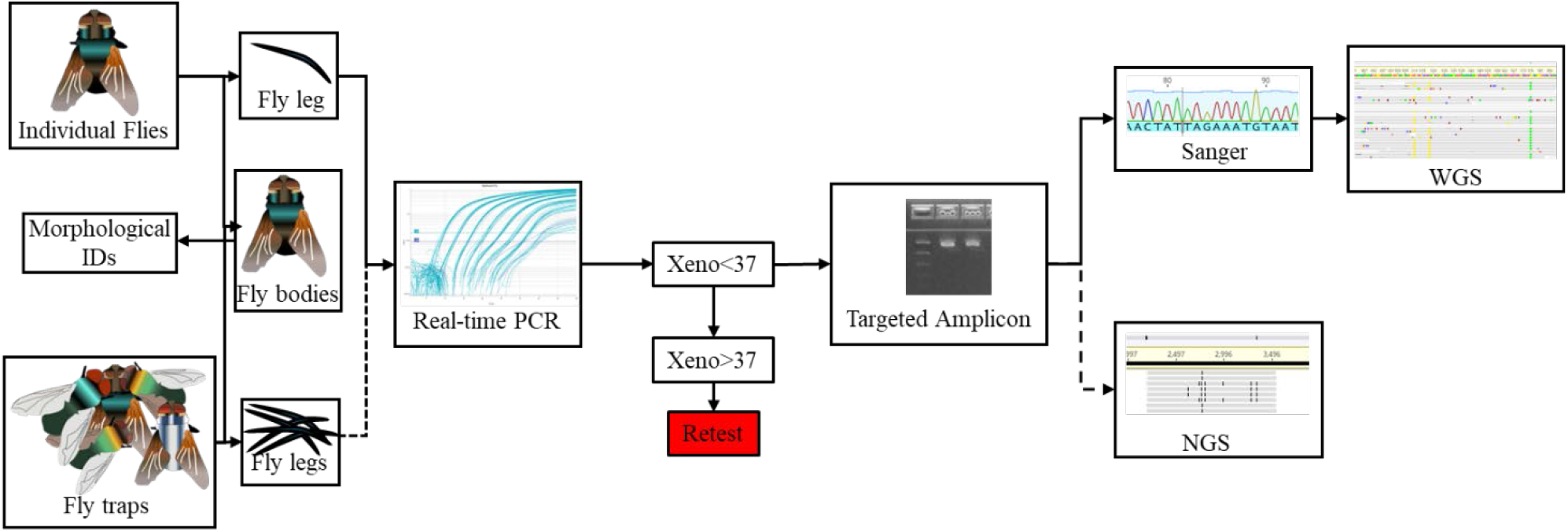
Workflow for identification of *Coc. hominivorax*. Sample types include individual flies or maggots, or bulk samples from a fly trap. Once the samples are analyzed by real-time PCR, they will be used to amplify regions of interest; and those amplicons will be sequenced by either Sanger (individual flies or maggots) or by NGS (bulk fly trap samples).

Currently, sequencing approaches focus on the COI-5P region of the COXI mitochondrial gene, a region that is flanked by sequences common among many different insect species^19,22,23^. The differential power is derived from the analysis of Sanger sequences following amplification. This approach is sufficient for species-level identification but is slow and cumbersome, requiring specialized equipment if, for example, a mixed-species bulk fly sample is being assessed. The BOLD database compiles information from numerous laboratories on the COI-5P region of *Coc. hominivorax* and related species^24^. This database also includes primers used to amplify across Diptera species. Our sanger assays amplify the COI-5P region as well along with additional sequences surrounding this region, allowing for comparison against this sequence database.

In this manuscript, we assessed three different extraction techniques, MagMAX CORE kit, the Gentra Purgene tissue kit, and the Qiagen DNeasy kit within both the real-time PCR and Sanger assays. All kits worked sufficiently for each of the assays and produced high quality DNA for identification. The MagMAX CORE kit is adaptable to high-throughput analyses using the KingFisher Flex and therefore can be used for instances of high-throughput testing. Samples extracted using the commonly used Gentra Puregene and Qiagen DNeasy kits amplified the expected targets and remain as a backup for repeats and retests if the case arises.

To utilize the most current reference sequences, we attempted to sequence the whole genome of 29 maggots or flies. The results from this work yielded >90% complete sequences from different countries that are publicly available in the SRA database and referenced in Supplementary Table 1. WGS and Sanger sequences yielded identical results confirming the species identity of the samples utilized. In this analysis, we also provide a dataset covering 8 different countries with WGS references. Additional annotations will be uploaded when appropriate information becomes available.

The results of this manuscript provide a workflow for molecular detection and species-level identification of *Coc. hominivorax*. This includes two highly specific and highly sensitive real-time PCR assays targeting the rDNA region of the genome that differentiate *Coc. hominivorax* from other, closely related species. Additionally, four primer sets amplifying mitochondrial genomic regions were assessed in this paper to confirm the real-time PCR results and classify the *Coc. hominivorax* sequences loosely by geographic region, whereby we identified two distinct clades of *Coc. hominivorax* across the eight countries assessed. This workflow is rapid, scalable, and sensitive; and we expect ongoing optimization and refinement to make these tools indispensable for fully characterizing and monitoring the current outbreak situation for NWS.

## Acknowledgements

We would like to thank Todd Gilligan, Luke Tembrock, and Frida Zink for their guidance on insect field collection and available assays, platforms, and sequences. We would like to thank Kimberly Lohmeyer, Paul Hickner, Sohath Yusseff-Vanegas, and Mackenzie Tietjen for providing whole genome sequencing guidance, sequences, and a subset of non-target flies. We would like to thank Tyler Pohlenz for input regarding molecular detection and background for insect identification. Nianet Carrasquillo, Maria Romano, and Martha Chavez coordinated the procurement of samples from Central America and Mexico.

