## Supplemental Tables 1-5; Supplemental Figure 1 for "Validation of a molecular workflow for *Cochliomyia hominivorax* (New World screwworm) identification in field samples"

**Supplemental Table 1.** Previously Published Primers

| **Torres 2004** | |
|---|---|
| Locus | Sequence |
| CH01 | GCATTATACAGACTCAAACAC |
| | GTTTGTCTGGTTATAAGCTT |
| CH02 | TCTTATAGCGCATAGACACG |
| | CATGAAGGCTGCTTGTATG |
| CH05 | CAATCAACACAAACTCACG |
| | CACCGTTAGTATCGCATG |
| CH09 | ACTATACCGGCCTCACAC |
| | TTGTTGTTGTATGAGTGGTG |
| CH10 | TGTGGCAACATTTCTTTTG |
| | TTGGCAAAAAATGTCTCTG |
| CH11 | TTGCTGCAACATACTGGG |
| | TAATTTGTTGGCGGCTTC |
| Ch12 | ATGTCCATCACTTCAAGCAC |
| | TTTGCAGAGGAGTGAGCG |
| CH14 | TGCTGTTGGGTGTTGTAG |
| | TTATGATGACCTTCTCTACG |
| CH15 | ATCACAAGAGTAAACATTTC |
| | TGTGTGTGTAAGACATGC |
| CH20 | ATCCATGTCCTACTCAGC |
| | TGCATTTCAATTGCTTTC |

| **Taylor et al., 1996** | |
|---|---|
| Sp2 | CAGCTACTTTATGAGCTTTAGG |
| Sp3 | CATTTCAAGCTGTGTAAGCATC |
| C2F3 | GGTCAATGTTCAGAAATTTGTGG |
| Sp4 | GAGACCATTACTTGCTTTCAGTCATCT |
| N4R2 | GTAGGAGGAGCTGCTATATTAG |
| N4F3 | GCTTATTCATCGGTTGCTCA |
| P6 | ACATGAATTGGAGCTCGACCAGT |
| P7 | GGTACATTACCTCGGTTTCGTTATGAT |
| P4 | GGTCCCTTACGAATTGAATATATCCT |
| P5 | GAGTTCAAACCGGCGTAAGCCAGGT |

| **Narang&Degrugillier, 1995** | |
|---|---|
| 12sai | A CTA GGA TTA GAT ACC CTA TTA T |
| 16sa | ATG TTT TTG ATAAAC AGGCG |
| 16sar | CGCCTGTTTATCAAAAACAT |
| 16sbr | CTCCGGTTTGAACTCAGATC |
| t0ISO | ATTTACCCTATCAAGGTAA |
| 12sifi | CGGGCGATGTGTACATAATT |
| t-ISO | ATTTACCCTATCAAGGTAA |
| 12sair | AGGGTATCTAATCCTAGTTT |
| TTGATTTTTTGGTCATCCAGAAGT | |
| GATCAATATCATTGACC | Hall and Smith 1991 |

| **Mcdonagh, 2009** | |
|---|---|
| EF1aF | CCCATYTCCGGHTGGCACGG |
| EF1aR | GTCTCATGTCACGDACRGCG |

| **Taylor et al., 1996** | |
|---|---|
| Name | Sequence |
| SP1 | TACAATTTATCGCCTAAACTTCAGCC |
| SP4 | GAGACCATTACTTGCTTTCAGTCATCT |
| C2F3 | GGTCAATGTTCAGAAATTTGTGG |
| N3R2 | TGATTTCATTCATGATATAGTCC |
| N3F1 | CCTTTTGAATGTGGATTTGATCC |
| N5F3 | TTGTCTACTTTAAGACAATTAGG |
| P6 | ACATGAATTGGAGCTCGACCAGT |
| P5 | GAGTTCAAACCGGCGTAAGCCAGGT |
| Sp2 | CAGCTACTTTATGAGCTTTAGG |
| SP3 | CATTTCAAGCTGTGTAAGCATC |
| C3F1 | ACAGGTGCTATTGGAGCTAT |
| C3R1 | CCATAAACTGAATCAGCAAT |
| P6 | |
| P7 | GGTACATTACCTCGGTTTCGTTATGAT |
| P4 | GGTCCCTTACGAATTTGAATATATCCT |
| P5 | |

| **Simon et al., 1994** | |
|---|---|
| Name | Sequence |
| TI-N-8 | CTATCAAGGTAACCCTTTTTATCAGGCA |
| TI-N-4 | ATTTACCCTATCAAGGTAA |
| TM-J-206 | GCTAATTAAGCTACTGGGTTCAT |
| TM-N-193 | TGGGGTATGAACCCAGTAGC |
| N2-N-732 | GAAGTTTGGTTTAAACCTCC |
| N2-J-748 | ATTGGAGGTTTAAACCAAAC |
| N2-J-1006 | TAGGTGGATTACCTCCATTTTTAGG |
| TY-J-1460 | TACAATCTATCGCCTAAACTTCAGCC |
| C1-N-1560 | TGTTCCTACTATTCCGGCTCA |
| C1-J-1718 | GGGGGGTTTGGAAATTGATTAGTGCC |
| C1-J-1751 | GGAGCTCCTGACATAGCATTCCC |
| C1-J-1859 | GGTACAGGTTGAACTGTTTACCCTCC |
| C1-J-2183 | CAACATTTATTTTGATTTTTTGG |
| C1-N-2191 | CCCGGTAAAATTAAAATATAAACTTC |
| C1-J-2195 | TTGATTTTTTGGTCACCCTGAAGT |
| C1-N-2329 | ACTGTAAATATATGATGAGCTCA |
| C1-J-2441 | CCTACAGGAATTAAAATTTTTAGATGATTAGC |
| C1-N-2659 | GTCAATCCAGTAAATAATGG |
| C1-J-2797 | CCTCGACGTTATTCAGATTACC |
| TL2-N-3014 | TCCATTGCACTAATCTGCCATATTA |
| TL2-J-3037 | ATGGCAGATTAGTGCAATGG |
| C2-J-3138 | AGAGCTTCTCCTTTAATGGAACA |
| C2-J-3279 | GGACAACTTATTGAAATAATTTGAAC |
| C2-N-3389 | TCATAACTTCAGTATCATTG |
| C2-J-3400 | ATTGGTCATCAATGATACTGA |
| C2-N-3494 | GGTAAAATTACTCGATTATCAAC |
| C2-J-3571 | AGATGTAATTCATTCTTGA |
| C2-N-3661 | CCGCAAATTTCTGAACATTGACCA |
| C2-J-3696 | GAAATTTGCGGGGCTAATCATAG |

| | |
|---|---|
| Nest | GGTGGYATCGGHACAGTACC |
| Nest | CAGCRAMGTACCCACGACGC |
| NEW COIF | ATAGACGTTGAYACWCGAGC |
| NEW COIR | ATCCATTGCACTAATCTGCC |
| Nest | GTATTTGCTATTATRGCCGG |
| Nest | AATCCTAARAAATGTTGGGG |

| **Otranto et al. 2005** | |
|---|---|
| C2-J-2369 | Simon |
| TL2-N-3014 | Simon |
| GaF | GTAAACATAACATTCTTCCCCCAGCA |
| GaR | AAAAATGCTGGGGGAAGAATGTTA |
| LR-J-12887 | Simon |
| LR-N-13398 | Simon |
| D1 | CCCCCTGAATTTAAGCATAT |
| D2 | GTTAGACTCCTTGGTCCGTG |
| DGp | TGAAACTGCTTAGAATTAAGC |

| **Lessinger and Azeredo-Espin 2000** | |
|---|---|
| TM-N-193 | Simon |
| SR-J-14233 | Simon |
| TI-N-24 | Simon |
| SR-J-14776 | GCTGGCACGAATTTTGTC |

| **Litjens 2001** | |
|---|---|
| C1-J-2195 | Simon |
| L2-N-3014 | Simon |
| TM-N-193 | Simon |
| SR-J-14233 | Simon |
| ATP8-N-3931 | AAT TGG TGC TAT TTG AGG |
| TW-J-1287 | ACT AAT AGC CTT CAA AGC |
| TV-J-14122 | TTC AAT GTA AAT GAA ATG |
| N2-N327 | Oliviera 2006 |

| **Oliveira 2006** | |
|---|---|
| SR-J-14941 | AGCCAAAATAAAACTTA |
| N2-N-327 | CTAAACCTATTCAAGCTCC |
| TI-J-42 | TTACCTTGATAGGGTAAAT |
| TI-N-21 | ATCCTATCAAGATAGCCC |
| N3-J-5945 | CATGAATGAAATCAAGG |
| N5-N-6554 | GGATGATCAGAATATTTTGG |
| N5-N-6647 | GGTTCAATATGATTTATACC |

| **Griffiths 2009 Microsattelites** | |
|---|---|
| CheX01 | TTTTATATTCTTGCTTCACA |
| | CAAATGAAATTGTTCAAAGT |
| ChEX02 | GCTTCAGATTCTTGTTCAAA |
| | ATCCGAAGATGGTCAAAAT |
| ChEX03 | GATCGGTTTATGTATTACCAAGA |
| | GCGATACTCAGTCGAGAAAA |
| ChEX04 | GATCCTCGAGTTGACCAATA |

| | |
|---|---|
| TK-N-3785 | GTTTAAGAGACCAGTACTTG |
| TD-N-3862 | TTTAGTTTGACATACTAATGTTAT |
| A8-N-3914 | TCATCTAATTGGTGCTATTTGTGG |
| C3-J-5014 | TTATTTATTTTATCAGAAGT |
| C3-N-5460 | TCAACAAAATGTCAGTATCA |
| N3-J-5952 | GAATGAAATCAAGGTATATTA |
| N5-N-6562 | GATCAAGGTTGGTCAGAA |
| N5-J-6579 | TTCTGACCAACCTTGATC |
| N4-N-8484 | GCTAATATAGCAGCTCCTCC |
| N4-J-8502 | GTTGGAGGAGCTGCTATATTAG |
| N4-N-8718 | GCATATTCATCAGTTGCTCA |
| N4-N-8944 | GGAGCTTCAACATGAGCTTT |
| CB-J-10612 | CCAATTAATATTTCAAGATGATGAAA |
| CB-J-10933 | TATGTTTTACCTTGAGGACAAATATC |
| CB-N-10920 | TCCTCAAAATGATATTTGTCCTCA |
| CB-J-11338 | CACATTCAACCAGAATGATATTT |
| CB-N-11328 | AGCAAATAAAAAATATCATTC |
| CB-N-11367 | ATAACTCCTCCTAATTTATTAGGAAT |
| CB-J-11545 | ACATGAATTGGAGCTCGACCAGT |
| TS1-N-11683 | AAATTCTATCTTATGTTTTCAAAAC |
| N1-N-11841 | GGTACATTACCTCGGTTTCGTTATGAT |
| N1-N-12051 | GATTTTGCTGAAGGTGAATCAGA |
| N1-N-12248 | AAGCTAATCTAACTTCATAAG |
| N1-J-12314 | TAGAATTAGAAGATCAACCAGC |
| N1-J-12533 | GCATCACAAAAAGGCTGAGG |
| N1-J-12585 | GGTCCCTTACGAATTTGAATATATCCT |
| N1-N-12595 | GTAGCTTTTTTAACTTTATTAGAACG |
| LR-N-12866 | ACATGATCTGAGTTCAAACCGG |
| LR-J-12887 | CCGGTTTGAACTCAGATCATGT |
| LR-N-12945 | GCGACCTCGATGTTGGATTAA |
| LR-N-13176 | ACGAGAAGACCCTATAAATC |
| LR-J-13331 | TGATTATGCTACCTTTGCACAGT |
| LR-J-13375 | TCAGTGGGCAGGTTAGAC |
| LR-J-13417 | ATGTTTTTGTTAAACAGGCG |
| LR-N-13398 | CGCCTGTTTAACAAAAACAT |
| SR-J-14233 | AAGAGCGACGGGCGATGTGT |
| SR-N-14275 | AAGGTGGATTTGGTAGTAAA |
| SR-J-14508 | TACAAAACAGGTTCCTCTG |
| SR-N-14522 | ACTTAAAAAAATTTGGCGGT |
| SR-N-14588 | AAACTAGGATTAGATACCCTATTAT |
| SR-J-14612 | AGGGTATCTAATCCTAGTTT |
| SR-N-14756 | GACCAAATTGGTGCCAGCAGT |
| SR-N-14922 | AAGTTTTATTTTGGCTTA |

| **Caravalho 2009** | |
|---|---|
| 7F1a | AGC TAA ATC CCG AAA CTA AAC |
| 7R4 | CTG TKG ARC CNT ATC AGA C |
| 7R3a | ATC CTT ATC ATT ATT TTC ACC C |
| 7F4a | CTG TKG ARC CNT ATC AGA C |
| E3F | ATG AAT TTC AAI GTY AGY YWI ITG GAG |
| 7R7 | TTG GTT ACA CTC TAA AAT AAA TC |

| | |
|---|---|
| | CCCCATATACCTGGAAGTAA |

| **Christen 2015** | |
|---|---|
| F | CGGCGATAGTGGATGTGG |
| CR92A1R | CGGCGATAGAATGGGAGATA |
| J1A2F | CGGCGATAGAATGGGAGATA |
| R | GCGATAGTGGATGTGGGATT |

| **Cardoso 2016** | |
|---|---|
| Cyp6g1 | GAGCCCGATAAATTCGAACC |
| | ATCATGCCAAAACGTTCACC |
| foraging | TGGTGGGGTTTACAAAATCG |
| | TTATCCCAGCCGGTAACATC |
| GD | CATTCGGTGGTGCTAAGG |
| | CCGGGTCCAATGAAACC |
| Jonah65aiv | ACTGCTGCTCATTGTACTGATG |
| | TTGCTATTCCAGCCAGAGTG |
| Malvolio | TGGGGTGTGGGTATTTTAGC |
| | TAAAACACGACGCCAACG |
| PGRP-SC2 | CGTGGCTGGAATGTTATGG |
| | TTGACCACGAGCAACAGC |
| RPS6-p70 | TCACCAAGACGTACACCAAG |
| | TGTACATCCATCATTTCGTC |
| Smooth | TTCTTATTGCCCGATCATACAC |
| | AATCTTGCTCGGTGGTTG |
| Foraging | TCCCAAATACGATAAGGATG |
| | CATCACGGAATTCTTCATTG |
| | ACTTGCTTGGTCATAGATCG |
| | TTTGTAAACCCCACCAATAG |
| GD | AAAGGCTTTGCCTACTGCTG |
| | TGGTACCGTGTTCGTTCTTG |
| | AACGAAGCCAACTACATGAGC |
| | CATCACGGTAAGTGGTGAAG |
| Jonah65aiv | GCCTGGTGTGGTGGTTC |
| | CGATACCAGTATTAGATTTAATCC |
| Malvolio | CGTAAATTATGGGCCTTCAC |
| | GAATGTTTGGCATCTTCRTAC |
| | GTTTTTGCTCATGGCATG |
| | CGGCCATATGTATCACAAG |
| RPS6-p70 | ATGGCTGACGTGASYG |
| | AAGCACCTAATGACCACC |
| | CGTGGTCATGGTAAAGC |
| | TGTACATCCATCATTTCGTC |

| **Thyssen 2005** | |
|---|---|
| C1-j-2183 | Simon paper |
| TL2-N-3014 | |
| Tl-N-4 | |
| CMEG-AR | Lessinger 2004 |

| **Lessinger 2004** | |
|---|---|
| CMEG-A | ATGATATTCTTAACTGGATT |

| **Cardoso 2014** | |
|---|---|
| Actin | GCCATGTATGTTGCCATCC |
| | CGACCAGCCAAATCCAAAC |
| a-tub | GATTTTGACCACTCACAC |
| | AACGATTCAAGTTGGTG |
| EF1 | ACAATCAGCTTGAGAGG |
| | ATTGATATTGCTTTGTGG |
| gapdh | GTCAGTGACACCCACTCCTC |
| | TTGATCAAGTCGATGACACG |
| gstD1 | AAAATCAACCTGCCGATCC |
| | GCCAATAAAGCTAAATCGGCTA |
| RPL13a | CGAAGAATTGAACCTCTC |
| | AAGTGGAATGGACCAC |
| Rp49 | GCACCAAGCACTTCATCC |
| | AGTGGGAAGCATGTGACG |
| RPS17 | TCGTGTAAGAACCAAGAC |
| | GCAGATACGCTTGTTG |
| RPS18 | TGACTTCCTCTAACTTGG |
| | ACCAGTAGTCTTGGTGTG |
| SdhA | AAGATGATTGGAAATGG |
| | AAGGGCATACCATAATTC |

| **Amat 2023** | |
|---|---|
| C1-J-2495 | CAGCTACTTTATGAGCTTTAGG |
| C1-N-2800 | CATTTCAAGCTGTGTAAGCATC |

| **Alamalakala 2009** | |
|---|---|
| EcoRiF | CTCGTAGACTGCGTACC |
| R | AATTGGTACGCAGTCTAC |
| MseIF | GACGATGAGTCCTGAG |
| R | TACTCAGGACTCAT |
| E | GAC TGC GTA CCA ATT C |
| M | GAT GAG TCC TGA GTA AC |
| E | GAC TGC GTA CCA ATT C NNN |
| M | GAT GAG TCC TGA GTA AC NN |
| E1 | GAC TGC GTA CCA ATT C AGG |
| E2 | GAC TGC GTA CCA ATT C ACA |
| E3 | GAC TGC GTA CCA ATT C ACT |
| M1 | GAT GAG TCC TGA GTA A CTA |
| M2 | GAT GAG TCC TGA GTA A CTT |
| M3 | GAT GAG TCC TGA GTA A CTG |
| M4 | GAT GAG TCC TGA GTA A CTC |
| M5 | GAT GAG TCC TGA GTA A CAG |
| M6 | GAT GAG TCC TGA GTA A CAC |
| M7 | GAT GAG TCC TGA GTA A CAA |
| M8 | GAT GAG TCC TGA GTA A CAT |

| **Perdormo 2020** | |
|---|---|
| N2-J-1480 | TACAATTTATCGCCTAAACTTCAGCC |
| C2-N-3793 | GAGACCATTACTTGCTTTCAGTCATCT |
| C2-J-3662 | GGTCAATGTTCAGAAATTTGTGG |

| | |
|---|---|
| SR-J-14776 | GCTGGCACGAATTTTGTC |
| TI-N-24 | ATTTACCCTATCAAGGTAA |
| CMEGAR | AATCCAGTTAAGAATATCAT |
| TI-N-24 | |
| SR-J-14776 | |
| N2-N-1128 | CGTAATAATTTAATATAAAAGC |
| SR-J-14776 | |
| TM-N-193 | TGGGGTATGAACCCAGTAGC |
| CMEGAR | |

| **Bergamo 2015** | |
|---|---|
| 7Foa | GGTATACCA TACGCCCAAC |
| RN2 | AACAGTAATCCCTCGTACG |
| 7FIn1 | ATTGTGTCTCCCTGCAAGTG |
| 7R1aN | CGTTTAGTTTCTGGAGCC |
| 7FIn2 | ACCATCGGTGAGTTGAGAG |
| 7R3a | ATCCTTATCATTATTTT CACCC |
| 380F | ACGGATT CGCG CTATTTAGA |
| 61R | GATGACGAGCTT GTTGG TGA |
| 1648R | CCACTCCGCCTTTAGTTTGA |
| 2099F | TGTGCCAAAGTTGTTTCTG |
| 1514R | GTAAG GGGTCCGTCAAAACA |
| 2385F | CCTCGATATACAGACCGATA |

| **Folmer 1994** | |
|---|---|
| LCO1490 | GGTCAACAAATCATAAAGATATTGG |
| HCO-2198 | TGATTTTTTGGTCACCCTGGAGTTTA |
| LCO1606 | TTAATTGGAGATGATCAAAT |
| LCO1495 | ACAAATCATAAAGATATTGG |
| LCO1828 | GGAGCTGGTACAGGTTGAAC |
| HCO1862 | TGAACTGTTTACCCTCCTTT |
| HCO2042 | TGATCAGTACTTATTACAGC |
| HCO2799 | CCACGACGGTACATAGATTATGCTG |
| HCO2911 | GAAGCTTTAGTGTGTCAACGAGGGG |
| HCO2192 | TGATTTTTTGGTCACCCTGA |
| HCO2798 | CAACATTTTTTAGGATTAGC |

| **Malgorn 1999** | |
|---|---|
| F1 | TTA TCA TTA CCA GTA TTA GC |
| R1 | AGC TGA AGT AAA GTA AGC T |
| F2 | CTA CTT TAT GAG CTT TAG G |
| R2 | TTC AAG TTG TGT AAG CAT C |

| **Newcomb 1997** | |
|---|---|
| 7F1a | AGCTAAATCCCGAAACTAAAC |

| **Singh 2012** | |
|---|---|
| Calcad207F | GAC CAT TTG TTT TGG CTG CT |
| Calcad908R | AAT CAA TGG ATT GGG GAG AA |
| Calcad627F | AGT TCG ATT GGT GTG CTG TG |
| Calcad1011R | CAG AAA GCT ATG GCC GAT TC |
| Oescado300F | AAC CCA CCG ATA AGA GAC CA |

| | |
|---|---|
| N3-N-5942 | TGATTTCATTCATGATATAAGTCC |
| N5-J-7660 | TTCTGATCATCCCTGATC |
| N5-N-8853 | GTAAAATCTTATAATGCTGG |
| N5-J-8621 | GCTATAGCAGCTGGCAATCAAG |
| N4-N-9676 | CTTATGAACGGATGGGGAGTC |
| N4-J-9580 | TAGGAGGAGCAGCCATATTTGGC |
| NL-N-10740 | GAAGGTCTTGGGCTTTCT |
| NL-J-10725 | AAGCCCAAGACCTTC |
| CB-N-11872 | GATGCACCATTGGCATGT |

| **Yusseff-Vanegas 2016** | |
|---|---|
| LCO1490 | GGTCAACAAATCATAAAGATATTGG |
| CI-N-2776 | GGATAATCAGAATATCGTCGAGG |
| B1 | CCCATYTCCGGHTGGCACGG |
| C1 | CTCTCATGTCACGDACRGCG |
| D1.F | CCCCCTGAATTTAAGCATAT |
| D35.486.R | TCGGAAGGAACCAGCTACTA |
| ITS4 | TCCTCCGCTTATTGATATGC |
| ITS5.8 | GGGACGATGAAGAACGCAGC |

| **Fresia 2011** | |
|---|---|
| CRc-N-BVII-R | Duarte 2008 |
| SR-J-14875 | AAGTCACACAAAAATTTAC |
| TW-J-1287 | Lyra 2009 |
| C1-N-2320 | AATCCTAATAATCCAATAGC |
| TL2-J-3034 | |
| TK-N-3785 | Simon paper |

| **Duarte 2008** | |
|---|---|
| C1-N-1560 | |
| SR-J-14612 | |
| TM-N-193 | Simon Paper |
| Crc-J-BVII | TTTTATATTGAAGCGTCG |
| Crc-N-BVII R | TTGCACGGTTCAATATAA |
| CMegAR | |
| CMegAR | Lessinger 2004 |
| CRv-J-Cadu | TTGGTAAATCCTACTGT |
| SR-J-14941 | Oliviera 2006 |

| **McDonagh 2009b** | |
|---|---|
| COX1F1 | ATAGACGTTGAYACWCGAGC |
| COX1R1 | ATCCATTGCACTAATCTGCC |
| COX1IntF1 | GTATTTGCTATTATRGCCGG |
| CoX1 intR4 | AATCCTAARAAATGTTGGGG |
| B1 | CCCATYTCCGGHTGGCACGG |
| C1 | GTCTCATGTCACGDACRGCG |
| Fint | GGTGGYATCGGHACAGTACC |
| Rint | AGTTTCRACACGACCGACG |
| EF1 | ACAGCGACGGTTTGTCTCATGTC |
| EF2 | CACATTAACATTGTCGTGATTGG |
| EF3 | CCGATACCACCGATTTTGTA |
| EF4 | CCTGGTTCAAGGGATGGAA |

| | |
|---|---|
| Oescado594F | TCG ATA CTG TGG CTG GTG AA |
| Oescado770F | TTG AGA AAT CTG GGC AAA CC |
| Oescado1287R | TCC TTG GCT TCC GTT AAG AA |
| CadT2207F | AGT GTG GGC GAG GTA ATG TC |
| CadT2370F | CTG CCT TGA AAG CCA ACT TC |
| CadT3156R | CAG AAA GCT ATG GCC GAC TC |

| **Torres 2005** | |
|---|---|
| CH21 | GGCGAGCGAGTCCTATTAC |
| R | CAGTTCCAGGTCAATTTCC |
| CH23 | GTTAAGAATATCCTGCTAATG |
| R | TTTATATGTCCCATTGTGC |
| CH24 | CCCACCAAAAACAATGTC |
| R | GTTTCCCACCAGTTCAGG |
| CH25 | CACCTACCTTAAGTGACACG |
| R | TTTTTCCTCTTCTGACCTC |
| CH26 | TATTTTCCCTTCATTCTTG |
| R | GGTGTTTTAATACCCGTAAAC |
| CH28 | CAATCACCAACCAAAATG |
| R | TGTATAAAGATGCGATTGAC |
| CH29 | CATCTACGGTGGTGTAGG |
| R | ACATGATATTTTCGCTCTC |

| **Wells 2001** | |
|---|---|
| TY-J-1460 | TACAATTTATCGCCTAAACTTCAGCC |
| C1-N-1687 | CAATTTCCAAATCCTCCAATTAT |
| C1-J-1751 | GGATCACCTGATATAGCATTCCC |
| C1-N-1840 | AGGAGGATAAACAGTTCAC/TCC |
| C1-J-2183 | CAACATTTATTTTGATTTTTTGG |
| C1-N-2191 | CCCGGTAAAATTAAAATATAAACTTC |
| C1-J-2319 | TAGCTATTGGAC/TTATTAGG |
| C1-N-2293 | AGTAAACCAATTGCTAGTATAGC |
| C1-J-2495 | CAGCTACTTTATGAGCTTTAGG |
| C1-N-2514 | AACTCCAGTTAATCCTCCTAC |
| C1-N-2659 | GCTAATCCAGTGAATAATGG |
| C1-J-2792 | ATACCTCGACGTTATTCAGA |
| C1-N-2800 | CATTTCAAGT/CTGTGTAAGCATC |
| TL2-N-3014 | TCCAATGCACTAATCTGCCATATTA |
| C2-J-3138 | AGAGCCTCTCCTTTAATAGAACA |
| C2-N-3389 | TCATAAGTTCA[R]TATCATTG |
| C2-J-3408 | CAATGATAT/CTGAAGT/ATATGA |
| TK-N-3775 | GAGACCATTACTTGCTTTCAGTCATCT |

| **Sperling 1994** | |
|---|---|
| a | CAACATTTATTTTGATTTTTTGG |
| b | GAGACCATTACTTGCTTTCAGTCATCT |
| a | CAACATTTATTTTGATTTTTTGG |
| c | CTAATCCAGTGAATAATGG |

| **Friedrich and Tautz 1997** | |
|---|---|
| 687 | CGG TGG ATC ACT (C/T)GG |
| 427 | CCC(C/G)CGTAA(T/C)TTAAGCATAT |

| | |
|---|---|
| C1-J1751a | GGATCACCTGATATAGCATTCCC |
| C1-J-2183 | CAACATTTATTTTGATTTTTTGG |
| C1-J-2495 | CAGCTACTTTATGAGCTTTAGG |
| C1-N-1840 | AGGAGGATAAACAGTTCAC/TCC |
| C1-N-2191 | CCCGGTAAAATTAAAATATAAACTTC |
| C1-N-2659 | GCTAATCCAGTGAATAATGG |
| TL2-N-3013 | TCCATTACATATAATCTGCCATATTAG |
| TY-J-1460 | TACAATTTATCGCCTAAACTTCAGCC |
| UEA7 | TACAGTTGGAATAGACGTTGATAC |
| UEA10 | TCCAATGCACTAATCTGCCATATTA |
| D1.F | CCCCCTGAATTTAAGCATAT |
| D2.R | GTTAGACTCCTTGGTCCGTG |
| D1.R | CTCTCTATTCAGAGTTCTTTTC |
| D2.F | GAGGGAAAGTTGAAAAGAAC |
| D3 | GACCCGTCTTGAAACACGG |
| D7.R | CGACTTCCCTTACCTACAT |
| D3-5.R | TTACACACTCCTTAGCGGA |
| D35.486.R | TCGGAAGGAACCAGCTACTA |
| D35.742.F | TCTCAAACTTTAAATGG |
| D7.F | GACTGAAGTGGAGAAGGGT |
| SR-J-14233 | AAGAGCGACGGGCGATGTGT |
| SR-J-14776 | GCTGGCACGAATTTTGTC |
| TI-N-24 | ATTTACCCTATCAAGGTAA |
| TM-N-193 | TGGGGTATGAACCCAGTAGC |

| **Yusseff-Vanegas 2017** | |
|---|---|
| Primer 1 | TACAATTTATCGCCTAAACTTCAGCC |
| C1-N-2191 | CCCGGTAAAATTAAAATATAAACTTC |
| C1-J-1751 | GGAGCTCCTGACATAGCATTCCC |
| C2-N-3014 | TCCATTGCACTAATCTGCCATATTA |

| **Parsons 2000** | |
|---|---|
| L14816 | CCATCCAACATCTCAGCATGATGAAA |
| H15173 | CCCCTCAGAATGATATTTGTCCTCA |

| **Bogdanowicz 1993** | |
|---|---|
| S1751 | GGATCACCTGATATAGCATTCCC |
| S1859 | GGAACIGGATGAACWGTTTAYCCICC |
| S2792 | ATACCTCGACGTTATTCAGA |
| A2191 | CCCGGTAAAATTAAAATATAAACTTC |
| A3014 | TCCAATGCACTAATCTGCCATATTA |
| A3389 | TCATAAGTTCARTATCATTG |
| A3772 | GAGACCATTACTTGCTTTCAGTCATCT |

| **Lunt 1996** | |
|---|---|
| UEAl | GAATAATTCCCATAAATAGATTTACA |
| UEAZ | TCAAGATAAAGGAGGATAAACAGTTC |
| UEAl | TATAGCATTCCCACGAATAAATAA |
| UEA4 | AATTTCGGTCAGTTAATAATATAG |
| UEA5 | AGTTTTAGCAGGAGCAATTACTAT |

| | |
|---|---|
| 686 | CTCTCTATTCA(A/G)AGTTCTTT(G/C) |
| 427 | |
| 485 | ACCCGTCTTGAAACACGA |
| 689 | ACACACTCCTTAGCGGA |
| 476 | CTGAAGTGGAGAAGGGT |
| 477 | GACTTCCCTTACCTACAT |
| 486 | TCGGAAGGAACCAGCTACTA |
| 706 | GCCAGTTCTGCTTACC |
| 742 | CTCAAACTTTAAATGG |
| 487 | AGGGTTTCGTGTGAACAG |
| 703 | TTCCAAACC(A/C)TATCTC |
| 743 | CGATTTTCAAGGTCC |

| | |
|---|---|
| UEA6 | TTAATWCCWGTWGGNACNGCAATRATTAT |
| UEA7 | TACAGTTGGAATAGACGTTGATAC |
| UEA8 | AAAAATGTTGAGGGAAAAATGTTA |
| UEA9 | GTAAACCTAACATTTTTTCCTCAACA |
| UEA10 | TCCAATGCACTAATCTGCCATATTA |

| **Stevens and Wall 2001** | |
|---|---|
| D1F | CCCCCTGAATTTAAGCATAT |
| D1R | GTTAGACTCCTTGGTCCGTG |
| D1R | CTCTCTATTCAGAGTTCTTTTC |
| D2F | GAGGGAAAGTTGAAAAGAAC |
| D3-5F | GACCCGTCTTGAAACACGG |
| D7R | CGACTTCCCTTACCTACAT |
| D3-5R | TTACACACTCCTTAGCGGA |
| D3-5.486R | TCGGAAGGAACCAGCTACTA |
| D3-5.742F | TCTCAAACTTTAAATGG |
| D7F | GACTGAAGTGGAGAAGGGT |

| **Zhang 1997** | | |
|---|---|---|
| UEA1 | Ty-N-1438 | GAATAATTCCCATAAATAGAACA |
| UEA1d | TY-N-1438d | GAAWAATTCCYATAAWTARATTTACA |
| UEA2 | C1-N-1844 | TCAAGATAAAGGAGGATAAACAGTTC |
| UEA2d | C1-N-1843d | GMWARWGGWGGRTAWACWGTTCA |
| UEA3 | C1-J-1763 | TATAGCATTCCCACGAATAAATAA |
| UEA4 | C1-N-2087 | AATTTCGGTCAGTTAATAATATAG |
| UEA4d | C1-N-2096d | GANGTATTWARRTTTCGRTCWGTTA |
| UEA5 | C1-J-2090 | AGTTTTAGCAGGAGCAATTACTAT |
| UEA6 | C1-N-2395d | TTAATWCCWGTWGGNACNGCAATRATTA |
| UEA7 | C1-J-2369 | TACAGTTGGAATAGACGTTGATAC |
| UEA8 | C1-N-2735 | AAAAATGTTGAGGGAAAAATGTTA |
| UEA9 | C1-J-2753 | GTAAACCTAACATTTTTTCCTCAACA |
| UEA10 | TL2-N-3014 | TCCAATGCACTAATCTGCCATATTA |

### Supplemental Table 2. Sample Metadata

| Tube Number | Species | Sex | Accession | Collection Date | Collection State | Collection County |
|---|---|---|---|---|---|---|
| 1 | Lucilia sp. | F | 25P00371 | 11/24/25 | California | San Diego |
| 2 | Cochliomyia macellaria | F | 25P00371 | 11/24/25 | California | San Diego |
| 3 | Sarcophagidae | | 25P00374 | 11/24/25 | California | Imperial |
| 4 | Cochliomyia macellaria | M | 25P00374 | 11/24/25 | California | Imperial |
| 5 | Cochliomyia macellaria | F | 25P00374 | 11/24/25 | California | Imperial |
| 6 | Chrysomya megacephala | M | 25P00370 | 11/24/25 | Texas | Hidalgo |
| 7 | Chrysomya megacephala | F | 25P00370 | 11/24/25 | Texas | Hidalgo |
| 8 | Chrysomya rufifacies | F | 25P00370 | 11/24/25 | Texas | Hidalgo |
| 9 | Chrysomya rufifacies | F | 25P00370 | 11/24/25 | Texas | Hidalgo |
| 10 | Chrysomya megacephala | F | 25P00370 | 11/24/25 | Texas | Hidalgo |
| 11 | Cochliomyia hominivorax | F | NWS - COPEG | | | |
| 12 | Chrysomya megacephala | F | 25P00370 | 11/24/25 | Texas | Hidalgo |
| 13 | Cochliomyia macellaria | M | 25P00370 | 11/24/25 | Texas | Hidalgo |
| 14 | Chrysomya rufifacies | M | 25P00370 | 11/24/25 | Texas | Hidalgo |
| 15 | Cochliomyia macellaria | F | 25P00365 | 11/24/25 | Texas | Cameron |
| 16 | Chrysomya rufifacies | M | 25P00365 | 11/24/25 | Texas | Cameron |
| 17 | Cochliomyia hominivorax | M | NWS - COPEG | | | |
| 18 | Composomyiops callipes | F | ARS - Colony | | | |
| 19 | Cochliomyia macellaria | F | 25P00390 | 12/1/25 | Texas | Starr |
| 20 | | | 25P00390 | 12/1/25 | Texas | Starr |
| 21 | | | 25P00390 | 12/1/25 | Texas | Starr |
| 22 | Chrysomya rufifacies | F | 25P00390 | 12/1/25 | Texas | Starr |
| 23 | Chrysomya rufifacies | F | 25P00390 | 12/1/25 | Texas | Starr |
| 24 | Cochliomyia macellaria | F | 25P00390 | 12/1/25 | Texas | Starr |
| 25 | Chrysomya rufifacies | F | 25P00390 | 12/1/25 | Texas | Starr |
| 26 | | | 25P00390 | 12/1/25 | Texas | Starr |
| 27 | Cochliomyia macellaria | F | 25P00390 | 12/1/25 | Texas | Starr |
| 28 | Chrysomya megacephala | M | 25P00390 | 12/1/25 | Texas | Starr |
| 29 | Chrysomya rufifacies | M | 25P00390 | 12/1/25 | Texas | Starr |
| 30 | Cochliomyia macellaria | F | 25P00390 | 12/1/25 | Texas | Starr |
| 31 | Chrysomya rufifacies | F | 25P00390 | 12/1/25 | Texas | Starr |
| 32 | Cochliomyia macellaria | F | 25P00402 | 12/1/25 | Texas | Zapata |
| 33 | Cochliomyia macellaria | M | 25P00402 | 12/1/25 | Texas | Zapata |
| 34 | | | 25P00402 | 12/1/25 | Texas | Zapata |
| 35 | Cochliomyia macellaria | F | 25P00402 | 12/1/25 | Texas | Zapata |
| 36 | Chrysomya rufifacies | F | 25P00402 | 12/1/25 | Texas | Zapata |
| 37 | Cochliomyia macellaria | F | 25P00402 | 12/1/25 | Texas | Zapata |
| 38 | Chrysomya rufifacies | F | 25P00402 | 12/1/25 | Texas | Zapata |
| 39 | Cochliomyia macellaria | F | 25P00402 | 12/1/25 | Texas | Zapata |
| 40 | Cochliomyia macellaria | F | 25P00402 | 12/1/25 | Texas | Zapata |
| 41 | Cochliomyia macellaria | M | 25P00402 | 12/1/25 | Texas | Zapata |
| 42 | | | 25P00402 | 12/1/25 | Texas | Zapata |
| 43 | Cochliomyia macellaria | M | 25P00402 | 12/1/25 | Texas | Zapata |
| 44 | Chrysomya rufifacies | F | 25P00414 | 12/4/25 | Texas | Val Verde |
| 45 | Cochliomyia macellaria | M | 25P00414 | 12/4/25 | Texas | Val Verde |
| 46 | Chrysomya rufifacies | F | 25P00414 | 12/4/25 | Texas | Val Verde |
| 47 | Cochliomyia macellaria | M | 25P00414 | 12/4/25 | Texas | Val Verde |

| | | | | | | |
|---|---|---|---|---|---|---|
| 48 | | | 25P00414 | 12/4/25 | Texas | Val Verde |
| 49 | Cochliomyia macellaria | F | 25P00414 | 12/4/25 | Texas | Val Verde |
| 50 | Chrysomya rufifacies | M | 25P00414 | 12/4/25 | Texas | Val Verde |
| 51 | Cochliomyia macellaria | F | 25P00418 | 12/4/25 | Texas | Maverick/Webb |
| 52 | Cochliomyia macellaria | F | 25P00418 | 12/4/25 | Texas | Maverick/Webb |
| 53 | Cochliomyia macellaria | F | 25P00418 | 12/4/25 | Texas | Maverick/Webb |
| 54 | Phormia regina | M | 25P00418 | 12/4/25 | Texas | Maverick/Webb |
| 55 | Cochliomyia macellaria | F | 25P00418 | 12/4/25 | Texas | Maverick/Webb |
| 56 | Phormia regina | F | 25P00418 | 12/4/25 | Texas | Maverick/Webb |
| 57 | Cochliomyia macellaria | M | 25P00417 | 12/4/25 | Texas | Zapata |
| 58 | Cochliomyia macellaria | F | 25P00417 | 12/4/25 | Texas | Zapata |
| 59 | Chrysomya megacephala | M | 25P00417 | 12/4/25 | Texas | Zapata |
| 60 | Chrysomya rufifacies | M | 25P00419 | 12/4/25 | Arizona | Cochise |
| 61 | Chrysomya rufifacies | M | 25P00419 | 12/4/25 | Arizona | Cochise |
| 62 | Chrysomya rufifacies | M | 25P00419 | 12/4/25 | Arizona | Cochise |
| 63 | Chrysomya megacephala | F | 25P00419 | 12/4/25 | Arizona | Cochise |
| 64 | Cochliomyia macellaria | M | 25P00458 | 12/4/25 | California | San Diego |
| 65 | Cochliomyia macellaria | M | 25P00458 | 12/4/25 | California | San Diego |
| 66 | Cochliomyia macellaria | F | 25P00458 | 12/4/25 | California | San Diego |
| 67 | Cochliomyia macellaria | F | 25P00458 | 12/4/25 | California | San Diego |
| 68 | Cochliomyia macellaria | M | 25P00458 | 12/4/25 | California | San Diego |
| 69 | Cochliomyia macellaria | M | 25P00458 | 12/4/25 | California | San Diego |
| 70 | Cochliomyia macellaria | F | 25P00458 | 12/4/25 | California | San Diego |
| 71 | Cochliomyia macellaria | M | 25P00458 | 12/4/25 | California | San Diego |
| 72 | Cochliomyia macellaria | F | 25P00458 | 12/4/25 | California | San Diego |
| 73 | Cochliomyia macellaria | F | 25P00458 | 12/4/25 | California | San Diego |
| 74 | Cochliomyia macellaria | M | 25P00458 | 12/4/25 | California | San Diego |
| 75 | Cochliomyia macellaria | F | 25P00458 | 12/4/25 | California | San Diego |
| 76 | Cochliomyia macellaria | F | 25P00458 | 12/4/25 | California | San Diego |
| 77 | Cochliomyia macellaria | M | 25P00458 | 12/4/25 | California | San Diego |
| 78 | Cochliomyia hominivorax | | NWS - COPEG | | | |
| 79 | Cochliomyia macellaria | M | 25P00458 | 12/4/25 | California | San Diego |
| 80 | Cochliomyia macellaria | M | 25P00458 | 12/4/25 | California | San Diego |
| 81 | Cochliomyia macellaria | F | 25P00458 | 12/4/25 | California | San Diego |
| 82 | Cochliomyia macellaria | F | 25P00457 | 12/4/25 | Texas | Starr |
| 83 | Chrysomya rufifacies | F | 25P00457 | 12/4/25 | Texas | Starr |
| 84 | Chrysomya rufifacies | F | 25P00457 | 12/4/25 | Texas | Starr |
| 85 | | | 25P00457 | 12/4/25 | Texas | Starr |
| 86 | Chrysomya rufifacies | F | 25P00457 | 12/4/25 | Texas | Starr |
| 87 | Cochliomyia macellaria | F | 25P00457 | 12/4/25 | Texas | Starr |
| 88 | Chrysomya rufifacies | M | 25P00457 | 12/4/25 | Texas | Starr |
| 89 | Chrysomya rufifacies | F | 25P00457 | 12/4/25 | Texas | Starr |
| 90 | Cochliomyia macellaria | M | 25P00457 | 12/4/25 | Texas | Starr |
| 91 | | | 25P00457 | 12/4/25 | Texas | Starr |
| 92 | Cochliomyia macellaria | M | 25P00457 | 12/4/25 | Texas | Starr |
| 93 | Chrysomya rufifacies | M | 25P00457 | 12/4/25 | Texas | Starr |
| 94 | | | 25P00457 | 12/4/25 | Texas | Starr |
| 95 | Chrysomya rufifacies | F | 25P00457 | 12/4/25 | Texas | Starr |
| 96 | Cochliomyia hominivorax | | NWS - COPEG | | | |
| 97 | Cochliomyia macellaria | F | 25P00385 | 11/26/25 | Texas | Cameron |
| 98 | Cochliomyia macellaria | F | 25P00385 | 11/26/25 | Texas | Cameron |

| | | | | | | |
|---|---|---|---|---|---|---|
| 99 | Cochliomyia macellaria | M | 25P00385 | 11/26/25 | Texas | Cameron |
| 100 | Cochliomyia macellaria | F | 25P00385 | 11/26/25 | Texas | Cameron |
| 101 | Chrysomya rufifacies | F | 25P00385 | 11/26/25 | Texas | Cameron |
| 102 | Cochliomyia macellaria | F | 25P00385 | 11/26/25 | Texas | Cameron |
| 103 | | | 25P00385 | 11/26/25 | Texas | Cameron |
| 104 | Cochliomyia macellaria | F | 25P00385 | 11/26/25 | Texas | Cameron |
| 105 | Chrysomya rufifacies | F | 25P00385 | 11/26/25 | Texas | Cameron |
| 106 | Cochliomyia macellaria | F | 25P00385 | 11/26/25 | Texas | Cameron |
| 107 | Cochliomyia macellaria | M | 25P00385 | 11/26/25 | Texas | Cameron |
| 108 | Cochliomyia macellaria | F | 25P00385 | 11/26/25 | Texas | Cameron |
| 109 | Angioneur sp. | F | 25P00367 | 11/24/25 | Texas | Val Verde |
| 110 | Chrysomya rufifacies | M | 25P00367 | 11/24/25 | Texas | Val Verde |
| 111 | Chrysomya rufifacies | F | 25P00367 | 11/24/25 | Texas | Val Verde |
| 112 | | | 25P00367 | 11/24/25 | Texas | Val Verde |
| 113 | Chrysomya rufifacies | M | 25P00367 | 11/24/25 | Texas | Val Verde |
| 114 | Chrysomya rufifacies | M | 25P00367 | 11/24/25 | Texas | Val Verde |
| 115 | Cochliomyia macellaria | F | 25P00399 | 12/1/25 | California | Imperial |
| 116 | Cochliomyia macellaria | F | 25P00399 | 12/1/25 | California | Imperial |
| 117 | Cochliomyia macellaria | F | 25P00399 | 12/1/25 | California | Imperial |
| 118 | Cochliomyia hominivorax | M | NWS - COPEG | | | |
| 119 | Cochliomyia macellaria | F | 25P00399 | 12/1/25 | California | Imperial |
| 120 | Cochliomyia macellaria | M | 25P00399 | 12/1/25 | California | Imperial |
| 121 | Cochliomyia macellaria | F | 25P00399 | 12/1/25 | California | Imperial |
| 122 | Cochliomyia macellaria | F | 25P00399 | 12/1/25 | California | Imperial |
| 123 | Cochliomyia macellaria | M | 25P00399 | 12/1/25 | California | Imperial |
| 124 | Cochliomyia macellaria | F | 25P00399 | 12/1/25 | California | Imperial |
| 125 | Cochliomyia macellaria | F | 25P00399 | 12/1/25 | California | Imperial |
| 126 | Cochliomyia macellaria | F | 25P00399 | 12/1/25 | California | Imperial |
| 127 | Cochliomyia hominivorax | M | NWS - COPEG | | | |
| 128 | Cochliomyia macellaria | F | 25P00399 | 12/1/25 | California | Imperial |
| 129 | Chrysomya megacephala | M | 25P00393 | 12/1/25 | Texas | Hidalgo |
| 130 | Chrysomya megacephala | M | 25P00389 | 11/28/25 | Texas | Hidalgo |
| 131 | Chrysomya megacephala | M | 25P00389 | 11/28/25 | Texas | Hidalgo |
| 132 | Chrysomya rufifacies | M | 25P00405 | 12/1/25 | Texas | Val Verde |
| 133 | Chrysomya rufifacies | M | 25P00405 | 12/1/25 | Texas | Val Verde |
| 134 | Chrysomya rufifacies | M | 25P00405 | 12/1/25 | Texas | Val Verde |
| 135 | Chrysomya rufifacies | M | 25P00405 | 12/1/25 | Texas | Val Verde |
| 136 | Chrysomya rufifacies | M | 25P00405 | 12/1/25 | Texas | Val Verde |
| 137 | Chrysomya rufifacies | M | 25P00405 | 12/1/25 | Texas | Val Verde |
| 138 | Cochliomyia macellaria | M | 25P00405 | 12/1/25 | Texas | Val Verde |
| 139 | Chrysomya rufifacies | M | 25P00405 | 12/1/25 | Texas | Val Verde |
| 140 | Cochliomyia macellaria | F | 25P00405 | 12/1/25 | Texas | Val Verde |
| 141 | Phormia regina | F | 25P00405 | 12/1/25 | Texas | Val Verde |
| 142 | Cochliomyia macellaria | F | 25P00405 | 12/1/25 | Texas | Val Verde |
| 143 | Cochliomyia macellaria | M | 25P00405 | 12/1/25 | Texas | Val Verde |
| 144 | Cochliomyia macellaria | F | 25P00405 | 12/1/25 | Texas | Val Verde |
| 145 | Chrysomya rufifacies | M | 25P00404 | 12/2/25 | Arizona | Cochiese |
| 146 | Chrysomya rufifacies | M | 25P00404 | 12/2/25 | Arizona | Cochiese |
| 147 | Chrysomya rufifacies | F | 25P00404 | 12/2/25 | Arizona | Cochiese |
| 148 | Chrysomya megacephala | M | 25P00404 | 12/2/25 | Arizona | Cochiese |
| 149 | Chrysomya megacephala | M | 25P00404 | 12/2/25 | Arizona | Cochiese |

| | | | | | | |
|---|---|---|---|---|---|---|
| 150 | Chrysomya megacephala | M | 25P00404 | 12/2/25 | Arizona | Cochiese |
| 151 | Cochliomyia hominivorax | F | NWS - COPEG | | | |
| 152 | Cochliomyia macellaria | M | 25P00404 | 12/2/25 | Arizona | Cochise |
| 153 | Cochliomyia macellaria | M | 25P00404 | 12/2/25 | Arizona | Cochise |
| 154 | Chrysomya rufifacies | F | 25P00404 | 12/2/25 | Arizona | Cochise |
| 155 | Chrysomya rufifacies | M | 25P00404 | 12/2/25 | Arizona | Cochise |
| 156 | Chrysomya rufifacies | F | 25P00404 | 12/2/25 | Arizona | Cochise |
| 157 | Chrysomya megacephala | F | 25P00404 | 12/2/25 | Arizona | Cochise |
| 158 | Chrysomya rufifacies | M | 25P00455 | 12/4/25 | Texas | Webb |
| 159 | | | 25P00455 | 12/4/25 | Texas | Webb |
| 160 | Phormia regina | M | 25P00455 | 12/4/25 | Texas | Webb |
| 161 | Cochliomyia macellaria | M | 25P00455 | 12/4/25 | Texas | Webb |
| 162 | Cochliomyia macellaria | M | 25P00455 | 12/4/25 | Texas | Webb |
| 163 | Chrysomya rufifacies | M | 25P00455 | 12/4/25 | Texas | Webb |
| 164 | Chrysomya rufifacies | M | 25P00455 | 12/4/25 | Texas | Webb |
| 165 | | | 25P00455 | 12/4/25 | Texas | Webb |
| 166 | Chrysomya rufifacies | M | 25P00455 | 12/4/25 | Texas | Webb |
| 167 | Cochliomyia macellaria | F | 25P00455 | 12/4/25 | Texas | Webb |
| 168 | Cochliomyia macellaria | M | 25P00455 | 12/4/25 | Texas | Webb |
| 169 | Chrysomya megacephala | F | 25P00455 | 12/4/25 | Texas | Webb |
| 170 | | | 25P00455 | 12/4/25 | Texas | Webb |
| 171 | Cochliomyia macellaria | F | 25P00455 | 12/4/25 | Texas | Webb |
| 172 | Chrysomya megacephala | F | 25P00456 | 12/4/25 | Texas | Hidalgo |
| 173 | Cochliomyia macellaria | M | 25P00456 | 12/4/25 | Texas | Hidalgo |
| 174 | Sarcophagidae | F | 25P00456 | 12/4/25 | Texas | Hidalgo |
| 175 | Cochliomyia macellaria | M | 25P00456 | 12/4/25 | Texas | Hidalgo |
| 176 | Sarcophagidae | F | 25P00456 | 12/4/25 | Texas | Hidalgo |
| 177 | Chrysomya rufifacies | M | 25P00456 | 12/4/25 | Texas | Hidalgo |
| 178 | Chrysomya rufifacies | M | 25P00456 | 12/4/25 | Texas | Hidalgo |
| 179 | Chrysomya megacephala | M | 25P00456 | 12/4/25 | Texas | Hidalgo |
| 180 | | | 25P00456 | 12/4/25 | Texas | Hidalgo |
| 181 | Cochliomyia hominivorax | | NWS - COPEG | 12/4/25 | Texas | Hidalgo |
| 182 | | | 25P00456 | 12/4/25 | Texas | Hidalgo |
| 183 | Chrysomya rufifacies | F | 25P00456 | 12/4/25 | Texas | Hidalgo |
| 184 | Cochliomyia macellaria | M | 25P00456 | 12/4/25 | Texas | Hidalgo |
| 185 | Chrysomya rufifacies | M | 25P00456 | 12/4/25 | Texas | Hidalgo |
| 186 | Cochliomyia macellaria | M | 25P00456 | 12/4/25 | Texas | Hidalgo |
| 187 | Cochliomyia macellaria | F | 25P00456 | 12/4/25 | Texas | Hidalgo |
| 188 | Chrysomya megacephala | M | 25P00456 | 12/4/25 | Texas | Hidalgo |
| 189 | Cochliomyia macellaria | F | 25P00456 | 12/4/25 | Texas | Hidalgo |
| 190 | Cochliomyia macellaria | F | 25P00456 | 12/4/25 | Texas | Hidalgo |
| 191 | Cochliomyia macellaria | F | 25P00456 | 12/4/25 | Texas | Hidalgo |
| 192 | Cochliomyia macellaria | F | 25P00456 | 12/4/25 | Texas | Hidalgo |
| 193 | Cochliomyia macellaria | M | 25P00456 | 12/4/25 | Texas | Hidalgo |
| 194 | Cochliomyia macellaria | M | 25P00456 | 12/4/25 | Texas | Hidalgo |
| 195 | Cochliomyia macellaria | M | 25P00456 | 12/4/25 | Texas | Hidalgo |
| 196 | Cochliomyia macellaria | F | 25P00456 | 12/4/25 | Texas | Hidalgo |
| 197 | Chrysomya rufifacies | M | 25P00456 | 12/4/25 | Texas | Hidalgo |
| 198 | Chrysomya rufifacies | M | 25P00456 | 12/4/25 | Texas | Hidalgo |
| 199 | Chrysomya rufifacies | M | 25P00456 | 12/4/25 | Texas | Hidalgo |
| 200 | Chrysomya rufifacies | M | 25P00456 | 12/4/25 | Texas | Hidalgo |

| | | | | | | |
|---|---|---|---|---|---|---|
| 201 | Cochliomyia macellaria | F | 25P00456 | 12/4/25 | Texas | Hidalgo |
| 202 | Chrysomya megacephala | M | 25P00456 | 12/4/25 | Texas | Hidalgo |
| 203 | Chrysomya rufifacies | F | 25P00456 | 12/4/25 | Texas | Hidalgo |
| 204 | Cochliomyia macellaria | M | 25P00456 | 12/4/25 | Texas | Hidalgo |
| 205 | Chrysomya rufifacies | F | 25P00456 | 12/4/25 | Texas | Hidalgo |
| 206 | Chrysomya megacephala | M | 25P00456 | 12/4/25 | Texas | Hidalgo |
| 207 | Cochliomyia macellaria | M | 25P00456 | 12/4/25 | Texas | Hidalgo |
| 208 | Cochliomyia hominivorax | | NWS - COPEG | | | |
| 209 | Cochliomyia macellaria | F | 25P00458 | 12/4/25 | California | Arizona |
| 210 | Cochliomyia macellaria | M | 25P00458 | 12/4/25 | California | Arizona |
| 211 | Cochliomyia macellaria | M | 25P00458 | 12/4/25 | California | Arizona |
| 212 | Chrysomya rufifacies | M | 25P00458 | 12/4/25 | California | Arizona |
| 213 | Chrysomya rufifacies | M | 25P00458 | 12/4/25 | California | Arizona |
| 214 | | | 25P00458 | 12/4/25 | California | Arizona |
| 215 | Chrysomya rufifacies | M | 25P00458 | 12/4/25 | California | Arizona |
| 216 | Chrysomya rufifacies | M | 25P00458 | 12/4/25 | California | Arizona |
| 217 | Cochliomyia macellaria | M | 25P00458 | 12/4/25 | California | Arizona |
| 218 | Mix 1 | | | | | |
| 219 | Cochliomyia macellaria | F | 25P00458 | 12/4/25 | California | Arizona |
| 220 | Cochliomyia macellaria | F | 25P00458 | 12/4/25 | California | Arizona |
| 221 | Cochliomyia macellaria | F | 25P00458 | 12/4/25 | California | Arizona |
| 222 | Cochliomyia macellaria | M | 25P00458 | 12/4/25 | California | Arizona |
| 223 | Cochliomyia macellaria | M | 25P00458 | 12/4/25 | California | Arizona |
| 224 | Cochliomyia macellaria | M | 25P00458 | 12/4/25 | California | Arizona |
| 225 | Cochliomyia macellaria | F | 25P00458 | 12/4/25 | California | Arizona |
| 226 | Cochliomyia macellaria | M | 25P00458 | 12/4/25 | California | Arizona |
| 227 | Chrysomya rufifacies | F | 25P00458 | 12/4/25 | California | Arizona |
| 228 | Cochliomyia macellaria | M | 25P00458 | 12/4/25 | California | Arizona |
| 229 | Cochliomyia macellaria | F | 25P00458 | 12/4/25 | California | Arizona |
| 230 | Cochliomyia macellaria | F | 25P00458 | 12/4/25 | California | Arizona |
| 231 | Lucilia sp. | F | 25P00528 | 12/11/25 | New Mexico | Luna |
| 232 | Chrysomya rufifacies | F | 25P00528 | 12/11/25 | New Mexico | Luna |
| 233 | Cochliomyia macellaria | F | 25P00528 | 12/11/25 | New Mexico | Luna |
| 234 | Chrysomya rufifacies | M | 25P00541 | 12/15/25 | New Mexico | Luna |
| 235 | Chrysomya rufifacies | M | 25P00541 | 12/15/25 | New Mexico | Luna |
| 236 | Cochliomyia macellaria | F | 25P00529 | 12/11/25 | California | San Diego |
| 237 | Cochliomyia macellaria | M | 25P00529 | 12/11/25 | California | San Diego |
| 238 | Cochliomyia macellaria | M | 25P00529 | 12/11/25 | California | San Diego |
| 239 | Cochliomyia hominivorax | | NWS - COPEG | | | |
| 240 | | | | | | |
| 241 | Cochliomyia macellaria | M | 25P00529 | 12/11/25 | California | San Diego |
| 242 | Cochliomyia macellaria | F | 25P00529 | 12/11/25 | California | San Diego |
| 243 | Cochliomyia macellaria | M | 25P00529 | 12/11/25 | California | San Diego |
| 244 | | | | | | |
| 245 | Sarcophagidae | M | 25P00527 | 12/11/25 | Arizona | Cochise |
| 246 | | | 25P00527 | 12/11/25 | Arizona | Cochise |
| 247 | Chrysomya rufifacies | M | 25P00527 | 12/11/25 | Arizona | Cochise |
| 248 | Cochliomyia macellaria | F | 25P00539 | 12/11/25 | California | Imperial |
| 249 | Cochliomyia macellaria | F | 25P00539 | 12/11/25 | California | Imperial |
| 250 | Mix 2 | | | | | |
| 251 | Cochliomyia macellaria | F | 25P00539 | 12/11/25 | California | Imperial |

| | | | | | | |
|---|---|---|---|---|---|---|
| 252 | Chrysomya rufifacies | F | 25P00539 | 12/11/25 | California | Imperial |
| 253 | Cochliomyia macellaria | F | 25P00525 | 12/8/25 | California | San Diego |
| 254 | Cochliomyia macellaria | F | 25P00525 | 12/8/25 | California | San Diego |
| 255 | Cochliomyia macellaria | F | 25P00525 | 12/8/25 | California | San Diego |
| 256 | Cochliomyia macellaria | F | 25P00525 | 12/8/25 | California | San Diego |
| 257 | Cochliomyia macellaria | F | 25P00525 | 12/8/25 | California | San Diego |
| 258 | Cochliomyia macellaria | F | 25P00525 | 12/8/25 | California | San Diego |
| 259 | Cochliomyia macellaria | F | 25P00554 | 12/15/25 | California | Imperial |
| 260 | Cochliomyia macellaria | F | 25P00554 | 12/15/25 | California | Imperial |
| 261 | Cochliomyia macellaria | F | 25P00554 | 12/15/25 | California | Imperial |
| 262 | Cochliomyia macellaria | F | 25P00554 | 12/15/25 | California | Imperial |
| 263 | Cochliomyia macellaria | F | 25P00554 | 12/15/25 | California | Imperial |
| 264 | Cochliomyia macellaria | M | 25P00554 | 12/15/25 | California | Imperial |
| 265 | Cochliomyia macellaria | F | 25P00554 | 12/15/25 | California | Imperial |
| 266 | Cochliomyia macellaria | M | 25P00554 | 12/15/25 | California | Imperial |
| 267 | Cochliomyia macellaria | F | 25P00554 | 12/15/25 | California | Imperial |
| 268 | Cochliomyia macellaria | F | 25P00554 | 12/15/25 | California | Imperial |
| 269 | Cochliomyia macellaria | F | 25P00491 | 12/8/25 | Texas | Cameron |
| 270 | Cochliomyia macellaria | F | 25P00491 | 12/8/25 | Texas | Cameron |
| 271 | Cochliomyia macellaria | M | 25P00491 | 12/8/25 | Texas | Cameron |
| 272 | Chrysomya megacephala | M | 25P00491 | 12/8/25 | Texas | Cameron |
| 273 | Chrysomya rufifacies | M | 25P00491 | 12/8/25 | Texas | Cameron |
| 274 | Sarcophagidae | | 25P00491 | 12/8/25 | Texas | Cameron |
| 275 | Cochliomyia macellaria | M | 25P00491 | 12/8/25 | Texas | Cameron |
| 276 | Cochliomyia hominivorax | | 25P00491 | 12/8/25 | Texas | Cameron |
| 277 | Cochliomyia macellaria | M | 25P00491 | 12/8/25 | Texas | Cameron |
| 278 | Cochliomyia macellaria | M | 25P00491 | 12/8/25 | Texas | Cameron |
| 279 | Mix 3 | | | | | |
| 280 | Cochliomyia macellaria | M | 25P00491 | 12/8/25 | Texas | Cameron |
| 281 | Cochliomyia macellaria | F | 25P00484 | 12/8/25 | California | Imperial |
| 282 | Cochliomyia macellaria | F | 25P00484 | 12/8/25 | California | Imperial |
| 283 | | | 25P00484 | 12/8/25 | California | Imperial |
| 284 | Cochliomyia macellaria | F | 25P00484 | 12/8/25 | California | Imperial |
| 285 | Cochliomyia macellaria | F | 25P00484 | 12/8/25 | California | Imperial |
| 286 | Sarcophagidae | | 25P00484 | 12/8/25 | California | Imperial |
| 287 | Cochliomyia macellaria | F | 25P00484 | 12/8/25 | California | Imperial |
| 288 | Cochliomyia macellaria | F | 25P00484 | 12/8/25 | California | Imperial |
| 289 | Cochliomyia hominivorax | | | | | |
| 290 | Cochliomyia hominivorax | | | | | |
| 291 | Chrysomya rufifacies | F | 25P00458 | 12/4/25 | California | San Diego |
| 292 | Cochliomyia macellaria | M | 25P00539 | 12/11/25 | California | Imperial |
| 293 | Cochliomyia macellaria | F | 25P00491 | 12/8/25 | Texas | Cameron |
| 294 | Chrysomya megacephala | M | 25P00518 | * No collection date, received at NVSL 12/9/25 | Texas | Starr |
| 295 | Cochliomyia macellaria | F | 25P00518 | | Texas | Starr |
| 296 | Sarcophagidae | | 25P00518 | | Texas | Starr |
| 297 | Sarcophagidae | | 25P00518 | | Texas | Starr |
| 298 | Chrysomya rufifacies | M | 25P00518 | | Texas | Starr |
| 299 | Chrysomya rufifacies | F | 25P00518 | | Texas | Starr |
| 300 | Cochliomyia macellaria | F | 25P00490 | 12/8/25 | Texas | Maverick/Webb |

| | | | | | | |
|---|---|---|---|---|---|---|
| 301 | Mix 4 | | | | | |
| 302 | Phormia regina | M | 25P00490 | 12/8/25 | Texas | Maverick/Webb |
| 303 | Cochliomyia macellaria | M | 25P00490 | 12/8/25 | Texas | Maverick/Webb |

| **Count of Species** | **Sex** | | | |
|---|---|---|---|---|
| *Species* | *F* | *M* | *(blank)* | **Grand Total** |
| Cochliomyia macellaria | 95 | 60 | | 155 |
| Chrysomya rufifacies | 29 | 42 | | 71 |
| Chrysomya megacephala | 7 | 15 | | 22 |
| Cochliomyia hominivorax | 2 | 3 | 8 | 13 |
| Sarcophagidae | 2 | 1 | 5 | 8 |
| Phormia regina | 2 | 3 | | 5 |
| Lucilia sp. | 2 | | | 2 |
| Mix 1 | | | 1 | 1 |
| Angioneur sp. | 1 | | | 1 |
| Composomyiops callipes | 1 | | | 1 |
| Mix 2 | | | 1 | 1 |
| Mix 3 | | | 1 | 1 |
| Mix 4 | | | 1 | 1 |
| (blank) | | | | |
| Grand Total | 141 | 124 | 17 | 282 |

#### Whole genome sequences

| **Species** | **Isolate Name** | **Isolate Source** | **Date Collected** | **Collection Location** | **SRA Accession** |
|---|---|---|---|---|---|
| Cochliomyia hominivorax | Belize-Maggot-402025 | Not Collected | Not Collected | Belize | SRR37579083 |
| Cochliomyia hominivorax | Female-Fertile-NWS-fly-251212 | COPEG fertile specimen | 12/12/25 | Panama | SRR37579082 |
| Cochliomyia hominivorax | Female-Fertile-NWS-fly-251114 | COPEG fertile specimen | 11/14/25 | Panama | SRR37579071 |
| Cochliomyia hominivorax | Female-Fertile-NWS-fly-251121 | COPEG fertile specimen | 11/21/25 | Panama | SRR37579061 |
| Cochliomyia hominivorax | Female-Fertile-NWS-fly-251205 | COPEG fertile specimen | 12/5/25 | Panama | SRR37579060 |
| Cochliomyia hominivorax | Belize-Maggot-417 | Not Collected | Not Collected | Belize | SRR37579059 |
| Cochliomyia hominivorax | Male-Fertile-NWS-fly-251205 | COPEG fertile specimen | 12/5/25 | Panama | SRR37579058 |
| Cochliomyia hominivorax | Male-Fertile-NWS-fly-251114 | COPEG fertile specimen | 11/14/25 | Panama | SRR37579057 |
| Cochliomyia hominivorax | Panama-Maggot | Not Collected | Not Collected | Panama | SRR37579056 |
| Cochliomyia hominivorax | Female-Fertile-NWS-fly-251219 | Not Collected | 12/19/25 | Panama | SRR37579055 |
| Cochliomyia hominivorax | Female-Sterile-NWS-fly-251128 | COPEG sterile specimen | 11/28/25 | Panama | SRR37579081 |
| Cochliomyia hominivorax | Female-Fertile-NWS-fly-251128 | COPEG fertile specimen | 11/28/25 | Panama | SRR37579080 |
| Cochliomyia hominivorax | Female-Sterile-NWS-fly-251121 | COPEG sterile specimen | 12/21/25 | Panama | SRR37579079 |
| Cochliomyia hominivorax | Female-Sterile-NWS-fly-251212 | COPEG sterile specimen | 12/12/25 | Panama | SRR37579078 |
| Cochliomyia hominivorax | Male-Sterile-NWS-fly-251205 | COPEG sterile specimen | 12/5/25 | Panama | SRR37579077 |
| Cochliomyia hominivorax | Male-Fertile-NWS-fly-251128 | COPEG fertile specimen | 11/28/25 | Panama | SRR37579076 |
| Cochliomyia hominivorax | Male-Sterile-NWS-fly-251219 | COPEG sterile specimen | 12/19/25 | Panama | SRR37579075 |
| Cochliomyia hominivorax | Male-Fertile-NWS-fly-251212 | COPEG fertile specimen | 12/12/25 | Panama | SRR37579074 |
| Cochliomyia hominivorax | Male-Fertile-NWS-fly-251121 | COPEG fertile specimen | 11/21/25 | Panama | SRR37579073 |
| Cochliomyia hominivorax | Male-Sterile-NWS-fly-251128 | COPEG sterile specimen | 11/28/25 | Panama | SRR37579072 |
| Cochliomyia hominivorax | Male-Sterile-NWS-fly-251114 | COPEG sterile specimen | 11/14/25 | Panama | SRR37579070 |
| Cochliomyia hominivorax | Female-Sterile-NWS-fly-251219 | COPEG sterile specimen | 12/19/25 | Panama | SRR37579069 |
| Cochliomyia hominivorax | Female-Sterile-NWS-fly-251205 | COPEG sterile specimen | 12/5/25 | Panama | SRR37579068 |
| Cochliomyia hominivorax | Male-Sterile-NWS-fly-251212 | COPEG sterile specimen | 12/12/25 | Panama | SRR37579067 |
| Cochliomyia hominivorax | Male-Sterile-NWS-fly-251121 | COPEG sterile specimen | 11/21/25 | Panama | SRR37579066 |

| | | | | | |
|---|---|---|---|---|---|
| Cochliomyia hominivorax | Male-Fertile-NWS-fly-251219 | COPEG fertile specimen | 12/19/25 | Panama | SRR37579065 |
| Cochliomyia hominivorax | Female-Sterile-NWS-fly-251114 | COPEG sterile specimen | 11/14/25 | Panama | SRR37579064 |
| Cochliomyia hominivorax | Guatemala-Maggot | Not Collected | Not Collected | Guatemala | SRR37579063 |
| Cochliomyia hominivorax | Mexico-Maggot | Not Collected | Not Collected | Mexico | SRR37579062 |
| Cochliomyia macellaria | 1-macellaria-PCRFree | Not Collected | Not Collected | Not Collected | SRR37600761 |
| Cochliomyia macellaria | 1-macellaria | Not Collected | Not Collected | Not Collected | SRR37600760 |
| Chrysomya megacephala | 2-meqacephala-PCRFree | Not Collected | Not Collected | Not Collected | SRR37600759 |
| Chrysomya megacephala | 2-meqacephala | Not Collected | Not Collected | Not Collected | SRR37600758 |
| Chrysomya megacephala | 3-meqacephala-PCRFree | Not Collected | Not Collected | Not Collected | SRR37600757 |
| Chrysomya megacephala | 3-meqacephala | Not Collected | Not Collected | Not Collected | SRR37600756 |
| Chrysomya megacephala | 4-meqacephala-PCRFree | Not Collected | Not Collected | Not Collected | SRR37600755 |
| Chrysomya megacephala | 4-meqacephala | Not Collected | Not Collected | Not Collected | SRR37600754 |
| Phormia regina | 5-regina-PCRFree | Not Collected | Not Collected | Not Collected | SRR37600753 |
| Phormia regina | 5-regina | Not Collected | Not Collected | Not Collected | SRR37600752 |

**Sanger Amplicon Sequences**

| | | | | | |
|---|---|---|---|---|---|
| Cochliomyia hominivorax | Belize-Larva-1-Amplicon1 | Not Collected | Not Collected | Belize | SRR37599751 |
| Cochliomyia hominivorax | Belize-Larva-2-Amplicon1 | Not Collected | Not Collected | Belize | SRR37599750 |
| Cochliomyia hominivorax | Costa Rica-Larva-1-Amplicon1 | Not Collected | Not Collected | Costa Rica | SRR37599655 |
| Cochliomyia hominivorax | Costa Rica-Larva-2-Amplicon1 | Not Collected | Not Collected | Costa Rica | SRR37599644 |
| Cochliomyia hominivorax | Costa Rica-Larva-3-Amplicon1 | Not Collected | Not Collected | Costa Rica | SRR37599689 |
| Cochliomyia hominivorax | El Salvador-Larva-1-Amplicon1 | Not Collected | Not Collected | El Salvador | SRR37599678 |
| Cochliomyia hominivorax | El Salvador-Larva-2-Amplicon1 | Not Collected | Not Collected | El Salvador | SRR37599667 |
| Cochliomyia hominivorax | El Salvador-Larva-3-Amplicon1 | Not Collected | Not Collected | El Salvador | SRR37599740 |
| Cochliomyia hominivorax | Guatemala-Larva-1-Amplicon1 | Not Collected | Not Collected | Guatemala | SRR37599729 |
| Cochliomyia hominivorax | Guatemala-Larva-2-Amplicon1 | Not Collected | Not Collected | Guatemala | SRR37599634 |
| Cochliomyia hominivorax | Guatemala-Larva-3-Amplicon1 | Not Collected | Not Collected | Guatemala | SRR37599749 |
| Cochliomyia hominivorax | Honduras-Larva-1-Amplicon1 | Not Collected | Not Collected | Honduras | SRR37599710 |
| Cochliomyia hominivorax | Honduras-Larva-2-Amplicon1 | Not Collected | Not Collected | Honduras | SRR37599699 |
| Cochliomyia hominivorax | Honduras-Larva-3-Amplicon1 | Not Collected | Not Collected | Honduras | SRR37599662 |
| Cochliomyia hominivorax | Mexico-Larva-1-Amplicon1 | Not Collected | Not Collected | Mexico | SRR37599661 |
| Cochliomyia hominivorax | Mexico-Larva-2-Amplicon1 | Not Collected | Not Collected | Mexico | SRR37599660 |
| Cochliomyia hominivorax | Nicaragua-Larva-1-Amplicon1 | Not Collected | Not Collected | Nicaragua | SRR37599659 |
| Cochliomyia hominivorax | Nicaragua-Larva-2-Amplicon1 | Not Collected | Not Collected | Nicaragua | SRR37599658 |
| Cochliomyia hominivorax | Nicaragua-Larva-3-Amplicon1 | Not Collected | Not Collected | Nicaragua | SRR37599657 |
| Cochliomyia hominivorax | Panama-Larva-1-Amplicon1 | Not Collected | Not Collected | Panama | SRR37599656 |
| Cochliomyia hominivorax | Panama-Larva-2-Amplicon1 | Not Collected | Not Collected | Panama | SRR37599654 |
| Cochliomyia hominivorax | Panama-Larva-3-Amplicon1 | Not Collected | Not Collected | Panama | SRR37599653 |
| Cochliomyia hominivorax | COPEG-Larva-1-Amplicon1 | Not Collected | Not Collected | Panama | SRR37599652 |
| Cochliomyia hominivorax | COPEG-Larva-2-Amplicon1 | Not Collected | Not Collected | Panama | SRR37599651 |
| Cochliomyia hominivorax | COPEG-Larva-3-Amplicon1 | Not Collected | Not Collected | Panama | SRR37599650 |
| Cochliomyia hominivorax | COPEG-Larva-4-Amplicon1 | Not Collected | Not Collected | Panama | SRR37599649 |
| Cochliomyia hominivorax | COPEG-flyleg-11-Amplicon1 | Not Collected | Not Collected | Panama | SRR37599648 |
| Cochliomyia hominivorax | COPEG-flyleg-17-Amplicon1 | Not Collected | Not Collected | Panama | SRR37599647 |
| Cochliomyia hominivorax | COPEG-flyleg-96-Amplicon1 | Not Collected | Not Collected | Panama | SRR37599646 |
| Cochliomyia hominivorax | COPEG-flyleg-127-Amplicon1 | Not Collected | Not Collected | Panama | SRR37599645 |
| Cochliomyia hominivorax | COPEG-flyleg-151-Amplicon1 | Not Collected | Not Collected | Panama | SRR37599643 |
| Cochliomyia hominivorax | COPEG-flyleg-181-Amplicon1 | Not Collected | Not Collected | Panama | SRR37599642 |
| Cochliomyia hominivorax | Belize-Larva-1-Amplicon4 | Not Collected | Not Collected | Belize | SRR37599641 |
| Cochliomyia hominivorax | Belize-Larva-2-Amplicon4 | Not Collected | Not Collected | Belize | SRR37599640 |
| Cochliomyia hominivorax | Costa Rica-Larva-1-Amplicon4 | Not Collected | Not Collected | Costa Rica | SRR37599639 |

| | | | | | |
|---|---|---|---|---|---|
| Cochliomyia hominivorax | Costa Rica-Larva-2-Amplicon4 | Not Collected | Not Collected | Costa Rica | SRR37599638 |
| Cochliomyia hominivorax | Costa Rica-Larva-3-Amplicon4 | Not Collected | Not Collected | Costa Rica | SRR37599637 |
| Cochliomyia hominivorax | El Salvador-Larva-1-Amplicon4 | Not Collected | Not Collected | El Salvador | SRR37599692 |
| Cochliomyia hominivorax | El Salvador-Larva-2-Amplicon4 | Not Collected | Not Collected | El Salvador | SRR37599691 |
| Cochliomyia hominivorax | El Salvador-Larva-3-Amplicon4 | Not Collected | Not Collected | El Salvador | SRR37599690 |
| Cochliomyia hominivorax | Guatemala-Larva-1-Amplicon4 | Not Collected | Not Collected | Guatemala | SRR37599688 |
| Cochliomyia hominivorax | Guatemala-Larva-2-Amplicon4 | Not Collected | Not Collected | Guatemala | SRR37599687 |
| Cochliomyia hominivorax | Guatemala-Larva-3-Amplicon4 | Not Collected | Not Collected | Guatemala | SRR37599686 |
| Cochliomyia hominivorax | Honduras-Larva-1-Amplicon4 | Not Collected | Not Collected | Honduras | SRR37599685 |
| Cochliomyia hominivorax | Honduras-Larva-2-Amplicon4 | Not Collected | Not Collected | Honduras | SRR37599684 |
| Cochliomyia hominivorax | Honduras-Larva-3-Amplicon4 | Not Collected | Not Collected | Honduras | SRR37599683 |
| Cochliomyia hominivorax | Mexico-Larva-1-Amplicon4 | Not Collected | Not Collected | Mexico | SRR37599682 |
| Cochliomyia hominivorax | Mexico-Larva-2-Amplicon4 | Not Collected | Not Collected | Mexico | SRR37599681 |
| Cochliomyia hominivorax | Nicaragua-Larva-1-Amplicon4 | Not Collected | Not Collected | Nicaragua | SRR37599680 |
| Cochliomyia hominivorax | Nicaragua-Larva-2-Amplicon4 | Not Collected | Not Collected | Nicaragua | SRR37599679 |
| Cochliomyia hominivorax | Nicaragua-Larva-3-Amplicon4 | Not Collected | Not Collected | Nicaragua | SRR37599677 |
| Cochliomyia hominivorax | Panama-Larva-1-Amplicon4 | Not Collected | Not Collected | Panama | SRR37599676 |
| Cochliomyia hominivorax | Panama-Larva-2-Amplicon4 | Not Collected | Not Collected | Panama | SRR37599675 |
| Cochliomyia hominivorax | Panama-Larva-3-Amplicon4 | Not Collected | Not Collected | Panama | SRR37599674 |
| Cochliomyia hominivorax | COPEG-Larva-1-Amplicon4 | Not Collected | Not Collected | Panama | SRR37599673 |
| Cochliomyia hominivorax | COPEG-Larva-2-Amplicon4 | Not Collected | Not Collected | Panama | SRR37599672 |
| Cochliomyia hominivorax | COPEG-Larva-3-Amplicon4 | Not Collected | Not Collected | Panama | SRR37599671 |
| Cochliomyia hominivorax | COPEG-Larva-4-Amplicon4 | Not Collected | Not Collected | Panama | SRR37599670 |
| Cochliomyia hominivorax | COPEG-flyleg-11-Amplicon4 | Not Collected | Not Collected | Panama | SRR37599669 |
| Cochliomyia hominivorax | COPEG-flyleg-17-Amplicon4 | Not Collected | Not Collected | Panama | SRR37599668 |
| Cochliomyia hominivorax | COPEG-flyleg-96-Amplicon4 | Not Collected | Not Collected | Panama | SRR37599666 |
| Cochliomyia hominivorax | COPEG-flyleg-127-Amplicon4 | Not Collected | Not Collected | Panama | SRR37599665 |
| Cochliomyia hominivorax | COPEG-flyleg-151-Amplicon4 | Not Collected | Not Collected | Panama | SRR37599748 |
| Cochliomyia hominivorax | COPEG-flyleg-181-Amplicon4 | Not Collected | Not Collected | Panama | SRR37599747 |
| Cochliomyia hominivorax | Belize-Larva-1-Amplicon5 | Not Collected | Not Collected | Belize | SRR37599746 |
| Cochliomyia hominivorax | Belize-Larva-2-Amplicon5 | Not Collected | Not Collected | Belize | SRR37599745 |
| Cochliomyia hominivorax | Costa Rica-Larva-1-Amplicon5 | Not Collected | Not Collected | Costa Rica | SRR37599744 |
| Cochliomyia hominivorax | Costa Rica-Larva-2-Amplicon5 | Not Collected | Not Collected | Costa Rica | SRR37599743 |
| Cochliomyia hominivorax | Costa Rica-Larva-3-Amplicon5 | Not Collected | Not Collected | Costa Rica | SRR37599742 |
| Cochliomyia hominivorax | El Salvador-Larva-1-Amplicon5 | Not Collected | Not Collected | El Salvador | SRR37599741 |
| Cochliomyia hominivorax | El Salvador-Larva-2-Amplicon5 | Not Collected | Not Collected | El Salvador | SRR37599739 |
| Cochliomyia hominivorax | El Salvador-Larva-3-Amplicon5 | Not Collected | Not Collected | El Salvador | SRR37599738 |
| Cochliomyia hominivorax | Guatemala-Larva-1-Amplicon5 | Not Collected | Not Collected | Guatemala | SRR37599737 |
| Cochliomyia hominivorax | Guatemala-Larva-2-Amplicon5 | Not Collected | Not Collected | Guatemala | SRR37599736 |
| Cochliomyia hominivorax | Guatemala-Larva-3-Amplicon5 | Not Collected | Not Collected | Guatemala | SRR37599735 |
| Cochliomyia hominivorax | Honduras-Larva-1-Amplicon5 | Not Collected | Not Collected | Honduras | SRR37599734 |
| Cochliomyia hominivorax | Honduras-Larva-2-Amplicon5 | Not Collected | Not Collected | Honduras | SRR37599733 |
| Cochliomyia hominivorax | Honduras-Larva-3-Amplicon5 | Not Collected | Not Collected | Honduras | SRR37599732 |
| Cochliomyia hominivorax | Mexico-Larva-1-Amplicon5 | Not Collected | Not Collected | Mexico | SRR37599731 |
| Cochliomyia hominivorax | Mexico-Larva-2-Amplicon5 | Not Collected | Not Collected | Mexico | SRR37599730 |
| Cochliomyia hominivorax | Nicaragua-Larva-1-Amplicon5 | Not Collected | Not Collected | Nicaragua | SRR37599728 |
| Cochliomyia hominivorax | Nicaragua-Larva-2-Amplicon5 | Not Collected | Not Collected | Nicaragua | SRR37599727 |
| Cochliomyia hominivorax | Nicaragua-Larva-3-Amplicon5 | Not Collected | Not Collected | Nicaragua | SRR37599726 |
| Cochliomyia hominivorax | Panama-Larva-1-Amplicon5 | Not Collected | Not Collected | Panama | SRR37599725 |
| Cochliomyia hominivorax | Panama-Larva-2-Amplicon5 | Not Collected | Not Collected | Panama | SRR37599724 |
| Cochliomyia hominivorax | Panama-Larva-3-Amplicon5 | Not Collected | Not Collected | Panama | SRR37599723 |

| | | | | | |
|---|---|---|---|---|---|
| Cochliomyia hominivorax | COPEG-Larva-1-Amplicon5 | Not Collected | Not Collected | Panama | SRR37599722 |
| Cochliomyia hominivorax | COPEG-Larva-2-Amplicon5 | Not Collected | Not Collected | Panama | SRR37599721 |
| Cochliomyia hominivorax | COPEG-Larva-3-Amplicon5 | Not Collected | Not Collected | Panama | SRR37599636 |
| Cochliomyia hominivorax | COPEG-Larva-4-Amplicon5 | Not Collected | Not Collected | Panama | SRR37599635 |
| Cochliomyia hominivorax | COPEG-flyleg-11-Amplicon5 | Not Collected | Not Collected | Panama | SRR37599633 |
| Cochliomyia hominivorax | COPEG-flyleg-17-Amplicon5 | Not Collected | Not Collected | Panama | SRR37599632 |
| Cochliomyia hominivorax | COPEG-flyleg-96-Amplicon5 | Not Collected | Not Collected | Panama | SRR37599631 |
| Cochliomyia hominivorax | COPEG-flyleg-127-Amplicon5 | Not Collected | Not Collected | Panama | SRR37599630 |
| Cochliomyia hominivorax | COPEG-flyleg-151-Amplicon5 | Not Collected | Not Collected | Panama | SRR37599629 |
| Cochliomyia hominivorax | COPEG-flyleg-181-Amplicon5 | Not Collected | Not Collected | Panama | SRR37599628 |
| Cochliomyia hominivorax | Belize-Larva-1-Amplicon10 | Not Collected | Not Collected | Belize | SRR37599627 |
| Cochliomyia hominivorax | Belize-Larva-2-Amplicon10 | Not Collected | Not Collected | Belize | SRR37599626 |
| Cochliomyia hominivorax | Costa Rica-Larva-1-Amplicon10 | Not Collected | Not Collected | Costa Rica | SRR37599625 |
| Cochliomyia hominivorax | Costa Rica-Larva-2-Amplicon10 | Not Collected | Not Collected | Costa Rica | SRR37599624 |
| Cochliomyia hominivorax | Costa Rica-Larva-3-Amplicon10 | Not Collected | Not Collected | Costa Rica | SRR37599720 |
| Cochliomyia hominivorax | El Salvador-Larva-1-Amplicon10 | Not Collected | Not Collected | El Salvador | SRR37599719 |
| Cochliomyia hominivorax | El Salvador-Larva-2-Amplicon10 | Not Collected | Not Collected | El Salvador | SRR37599718 |
| Cochliomyia hominivorax | El Salvador-Larva-3-Amplicon10 | Not Collected | Not Collected | El Salvador | SRR37599717 |
| Cochliomyia hominivorax | Guatemala-Larva-1-Amplicon10 | Not Collected | Not Collected | Guatemala | SRR37599716 |
| Cochliomyia hominivorax | Guatemala-Larva-2-Amplicon10 | Not Collected | Not Collected | Guatemala | SRR37599715 |
| Cochliomyia hominivorax | Guatemala-Larva-3-Amplicon10 | Not Collected | Not Collected | Guatemala | SRR37599714 |
| Cochliomyia hominivorax | Honduras-Larva-1-Amplicon10 | Not Collected | Not Collected | Honduras | SRR37599713 |
| Cochliomyia hominivorax | Honduras-Larva-2-Amplicon10 | Not Collected | Not Collected | Honduras | SRR37599712 |
| Cochliomyia hominivorax | Honduras-Larva-3-Amplicon10 | Not Collected | Not Collected | Honduras | SRR37599711 |
| Cochliomyia hominivorax | Mexico-Larva-1-Amplicon10 | Not Collected | Not Collected | Mexico | SRR37599709 |
| Cochliomyia hominivorax | Mexico-Larva-2-Amplicon10 | Not Collected | Not Collected | Mexico | SRR37599708 |
| Cochliomyia hominivorax | Nicaragua-Larva-1-Amplicon10 | Not Collected | Not Collected | Nicaragua | SRR37599707 |
| Cochliomyia hominivorax | Nicaragua-Larva-2-Amplicon10 | Not Collected | Not Collected | Nicaragua | SRR37599706 |
| Cochliomyia hominivorax | Nicaragua-Larva-3-Amplicon10 | Not Collected | Not Collected | Nicaragua | SRR37599705 |
| Cochliomyia hominivorax | Panama-Larva-1-Amplicon10 | Not Collected | Not Collected | Panama | SRR37599704 |
| Cochliomyia hominivorax | Panama-Larva-2-Amplicon10 | Not Collected | Not Collected | Panama | SRR37599703 |
| Cochliomyia hominivorax | Panama-Larva-3-Amplicon10 | Not Collected | Not Collected | Panama | SRR37599702 |
| Cochliomyia hominivorax | COPEG-Larva-1-Amplicon10 | Not Collected | Not Collected | Panama | SRR37599701 |
| Cochliomyia hominivorax | COPEG-Larva-2-Amplicon10 | Not Collected | Not Collected | Panama | SRR37599700 |
| Cochliomyia hominivorax | COPEG-Larva-3-Amplicon10 | Not Collected | Not Collected | Panama | SRR37599698 |
| Cochliomyia hominivorax | COPEG-Larva-4-Amplicon10 | Not Collected | Not Collected | Panama | SRR37599697 |
| Cochliomyia hominivorax | COPEG-flyleg-11-Amplicon10 | Not Collected | Not Collected | Panama | SRR37599696 |
| Cochliomyia hominivorax | COPEG-flyleg-17-Amplicon10 | Not Collected | Not Collected | Panama | SRR37599695 |
| Cochliomyia hominivorax | COPEG-flyleg-96-Amplicon10 | Not Collected | Not Collected | Panama | SRR37599694 |
| Cochliomyia hominivorax | COPEG-flyleg-127-Amplicon10 | Not Collected | Not Collected | Panama | SRR37599693 |
| Cochliomyia hominivorax | COPEG-flyleg-151-Amplicon10 | Not Collected | Not Collected | Panama | SRR37599664 |
| Cochliomyia hominivorax | COPEG-flyleg-181-Amplicon10 | Not Collected | Not Collected | Panama | SRR37599663 |

**Supplemental Table 3.** Real-time PCR Master Mix

| Reagent | Initial Concentration (µM) | Final Concentration (nM) | Volume per Reaction (µl) |
|---|---|---|---|
| *Nuclease-free Water* | NA | NA | 10.25 |
| *TaqMan Fast Virus* | 4 | 1000 | 6.25 |
| *NWS Forward Primer* | 20 | 750 | 0.9375 |
| *NWS Reverse Primer* | 20 | 750 | 0.9375 |
| *Xeno Liz Probe* | 10 | 400 | 1 |
| *NSW Probe* | 10 | 250 | 0.625 |
| *$MgCL_2$* | 25 | 1000 | 1 |
| *Template* | NA | NA | 5 |
| | | ***Total Volume*** | 25 |

**Supplemental Table 4:** Real-time PCR Cycling Conditions

| Cycles | Block 1: NWS1 | Block 2: NWS4 | Block 3: NWS5 | Block 4: NWS10 | Block 5: NWS12 |
|---|---|---|---|---|---|
| **1** | 95°C \| 2min | -- | -- | -- | -- |
| **40** | 95°C \| 30sec<br>52°C \| 30sec<br>62°C \| 1:30min | 54°C \| 30sec<br>62°C \| 1min | 57°C \| 30sec<br>62°C \| 1:45min | 56°C \| 30sec<br>62°C \| 2:30min | 54°C \| 30sec<br>62°C \| 1:20min |
| **1** | 62°C \| 7min | -- | -- | -- | -- |

**Supplemental Table 5.** Sanger Master Mix

| Reagent | Volume (µl) | Final Concentration |
|---|---|---|
| *$H_2O$* | 11.55 | -- |
| *10xBFR* | 2.5 | 1X |
| *DMSO* | 1.25 | 5% |
| *$MgCl_2$* | 2.5 | 2.5nM |
| *dNTP* | 2 | 0.2nM |
| *Primers* | 2 | 0.4µM |
| *Taq* | 0.2 | 1U/rxn |
| *BSA* | 1 | 4% |
| *DNA* | 2 | -- |

**Supplemental Figure 1.** Primer Locations for real-time PCR and Sanger Primers

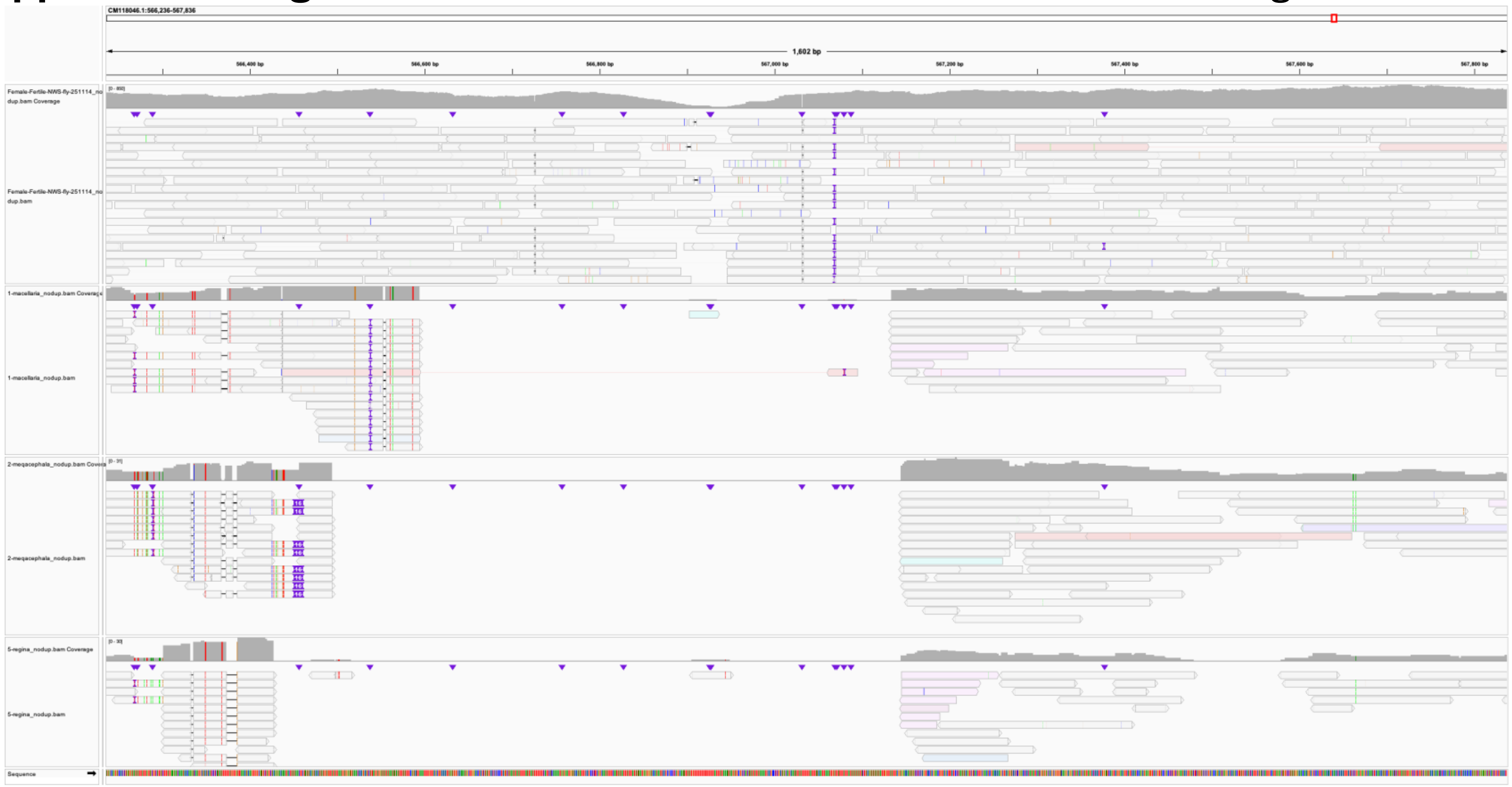

**A.** CM118046.1 – rDNA segment showing a gap in coverage for the non-NWS sequences

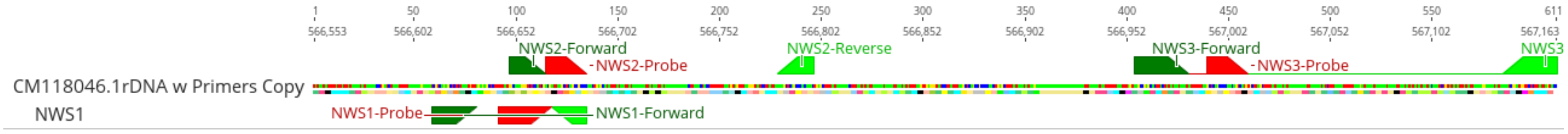

**B.** CM118046.1 – location of real-time PCR primers in this region.

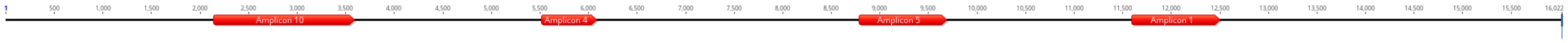

**C.** NC_002660 with Sanger amplicon locations.
